# High Resolution Ancestry Deconvolution for Next Generation Genomic Data

**DOI:** 10.1101/2021.09.19.460980

**Authors:** Helgi Hilmarsson, Arvind S. Kumar, Richa Rastogi, Carlos D. Bustamante, Daniel Mas Montserrat, Alexander G. Ioannidis

## Abstract

As genome-wide association studies and genetic risk prediction models are extended to globally diverse and admixed cohorts, ancestry deconvolution has become an increasingly important tool. Also known as local ancestry inference (LAI), this technique identifies the ancestry of each region of an individual’s genome, thus permitting downstream analyses to account for genetic effects that vary between ancestries. Since existing LAI methods were developed before the rise of massive, whole genome biobanks, they are computationally burdened by these large next generation datasets. Current LAI algorithms also fail to harness the potential of whole genome sequences, falling well short of the accuracy that such high variant densities can enable. Here we introduce Gnomix, a set of algorithms that address each of these points, achieving higher accuracy and swifter computational performance than any existing LAI method, while also enabling portable models that are particularly useful when training data are not shareable due to privacy or other restrictions. We demonstrate Gnomix (and its swift phase correction counterpart Gnofix) on worldwide whole-genome data from both humans and canids and utilize its high resolution accuracy to identify the location of ancient New World haplotypes in the Xoloitzcuintle, dating back over 100 generations. Code is available at https://github.com/AI-sandbox/gnomix.

## 1 Background & Summary

The importance of increasing inclusion in genome-wide association studies (GWAS) has been well documented^1^, but most worldwide populations are still highly underrepresented in large-scale genomic studies^2–4^, in part due to complexities involved in analyzing ancestrally diverse and admixed cohorts. Recently, advances have been made that utilize local ancestry inference (LAI) to help power diverse GWAS^5, 6^. These approaches place a burden on LAI algorithms to perform efficiently, reproducibly, and accurately on the increasingly large cohorts being studied.

Over the years several methods for local-ancestry inference have been published. SABER^7^, HAPAA^8^, and HAPMIX^9^ used Hidden Markov Models (HMMs). The accuracy and efficiency of these methods were later surpassed by LAMP^10^, a method that applied probability maximization within a sliding window, but could handle only a limited number of ancestries. Support Vector Machines (SVM)^11^ and Random Forests (RF) with a conditional random field (RFMix)^12^ were later explored for increased accuracy, with the latter currently considered to be state-of-the-art. Refinements including ELAI (HMMs)^13^, and Loter (dynamic programming with template matching and bagging) have been developed since^14^. Recently, LAI-Net, a local-ancestry estimation method based on a neural network, was demonstrated to provide competitive results^15^.

These past methods have been severely challenged by the ballooning numbers of samples in modern biobanks and the large number of sites in increasingly affordable whole genome sequences. As biobanks consisting of hundreds of thousands of individuals are now common^16–19^, and whole genome sequences of these biobanks are imminent, such limitations of existing LAI methods need to be addressed.

In this work, we present Gnomix, a flexible collection of new local ancestry algorithms capable of assigning ancestry labels to DNA segments from massive biobanks of either whole genome or single nucleotide polymorphism (SNP) array data with higher accuracy and swifter execution than any existing methods. The framework has two stages; a set of classifiers (base models) that perform an initial estimate of the ancestry probabilities within genomics windows, and a second stage consisting of another module (smoother) that learns to combine and refine these estimates, significantly increasing our accuracy. This framework is highly modular and supports a variety of combinations of classifier and smoother algorithms, enabling the user to select the configuration with a suitable trade-off between compute time, model size, and accuracy. We present an in-depth analysis of these configurations, and we recommend combinations that produce optimal trade-offs for the user according to their computational resources and their particular data.

For the base module’s classifiers (base models) we explore first the historically successful LAI classifiers (e.g. Random Forest and SVMs) followed by other classical machine learning classifiers (e.g. logistic regression and gradient boosting). In addition to these, we propose an innovation that generalizes the string kernel^20^. This method, which we name covariance reduction string kernel (CovRSK), adapts a sampling technique to reduce redundant features of the kernel and acts as a regularizer, improving the overall accuracy.

For the smoother we propose a data-driven approach, learning to capture the correlation structure of populations and the distribution of recombination breakpoints directly without relying on simplified models or assumptions. We explain how training the base models and smoother sequentially on the same data induces a covariate shift for this latter stage and present an alternative training pipeline which prevents this subtle bias. We compare various base models (both novel and existing) and show that XGBoost is the most successful smoother, surpassing alternatives like linear convolutional filters and conditional random fields (CRFs).

We show how several different instantiations of our method, Gnomix, widely outperform current state-of-the-art LAI methods in both accuracy and speed across a variety of datasets. We also introduce a novel phase correction algorithm, Gnofix, which uses a trained instance of a Gnomix smoother to phase segmented ancestry probabilities, such as the output of the base models, in a highly scalable manner. Finally, we demonstrate Gnomix-Gnofix capabilities by investigating the genomic history of New World dogs.

## 2 Results

### 2.1 Local ancestry inference with Gnomix

#### 2.1.1 System overview

We present a supervised method that uses haplotypes from a reference panel of single-ancestry individuals from differentiated population groups to learn a model that assigns an ancestry estimate to each segment of admixed chromosomes with high accuracy. We encode biallelic SNPs using the standard 0/1 format, zero for the reference allele and one for the alternative allele. For diploid organisms, such as humans, we consider maternal and paternal sequences independently and assume that they are correctly phased, although we can correct phasing errors with our new Gnofix algorithm, described below. If sequences are unphased, existing phasing algorithms can be employed before running Gnomix. Since we cannot have SNP-level ground-truth ancestry labels for admixed individuals, we simulate admixed progeny from single ancestry real genomes based on the human recombination map. We refer the reader to Section 3 for more details about dataset generation. Our proposed system splits each haplotype into windows of equal numbers of SNPs and outputs one ancestry estimate for each window. These initial probability estimates are then refined using the smoothing module, yielding the final ancestry predictions.

#### 2.1.2 Base module

In the first stage of the system, we produce initial ancestry estimates by assigning one model per window that learns to map the SNPs within that window to ancestry probabilities Figure 1a-d. The 0/1 encoded biallelic SNP sequences are segmented into W equally sized windows and each window is passed into a classifier. Because the learned classification function is not translation invariant (an identical sequence of zeros and ones might indicate a different ancestry depending on its location in the genome) a different classifier is applied at each windowed section of the input sequence. All window-specific classifiers have the same hyperparameter configuration, but each learns a different set of model parameters. Because neighbouring SNPs are usually inherited together (linkage), using a window-based approach that jointly processes co-located SNPs is a sensible approach. The independent treatment of windows results in highly parallelizable training and inference, leading to a significant time reduction that scales linearly with the number of available CPUs. Each model outputs a local ancestry probability estimate that can be thought of as an ancestry-aware mapping to translation-invariant features (the probability estimates) from the translation-variant SNP sequences. The former can be processed by convolutional or sliding-window based approaches (e.g. the smoother module).

**Figure 1.**
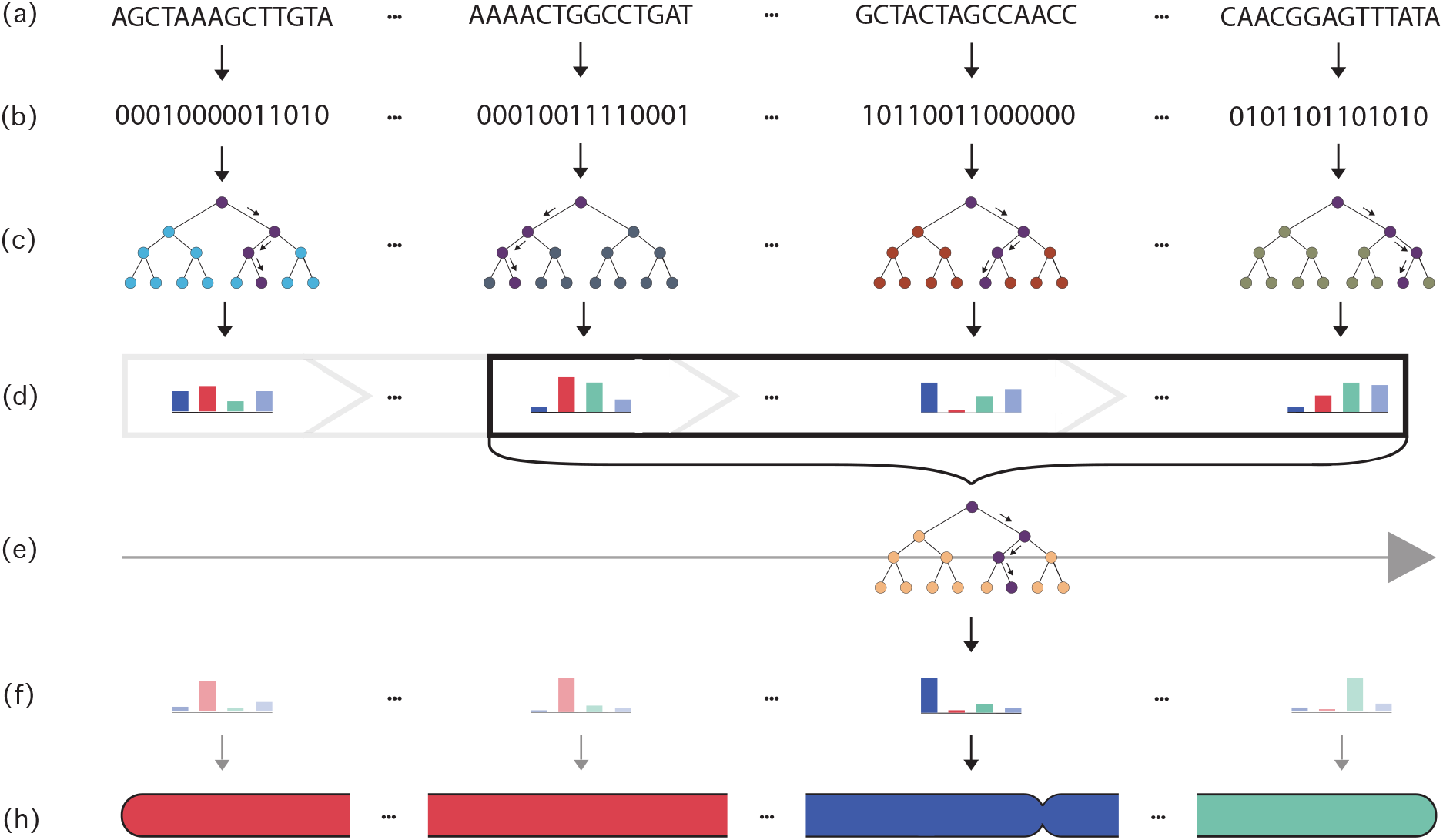
Schematic of the Gnomix algorithm. **a)** Genome sequence segmentation **b)** Binary (biallelic) encoding **c)** Local base classifiers’ learning procedure **d)** Probability estimates of local base classifiers **e-f)** Sliding smoother module is trained to learn to aggregate local base model probabilities **h)** Smoothing module outputs final local ancestry predictions along the chromosome.

Optimizing the window size of the classifiers results in a classic bias-variance trade-off. As the window size grows, each classifier sees more positions of the chromosome (more features) and provides more accuracy, *but* also yields a lower resolution series of genomic estimates, since larger windows necessarily span more of the genome. Such low resolution estimates can subsume small segments of ancestries that reflect old admixture events or can span ancestry switch-points, resulting in conflicting learning procedures as two different ancestries can then exist within the same window. On the other hand small windows give noisier probability estimates, leaving to the smoother the task of correcting the predictions. Furthermore, genotyping array data, which has a low density of SNPs, gives poor performance when small genetic windows are used, as a very low number of SNPs are available in each window. To alleviate these trade-offs we make use of overlapping windows, introducing a concept of context, *c*. With a non-negative context, each base-model sees *c* SNPs on either sides of its assigned segment. In this way given a window size of *w*, each base-model sees a total of *w* + 2*c* SNPs. This allows each model to see more SNPs without decreasing the prediction resolution along the genome.

Our proposed system is highly modular and enables a choice of many supervised discriminative classifier models for both the base and the smoothing module. This allows for easy replacements that are suited for a given task. For our system, we explored a broad spectrum of classifier families:

- Tree-based methods: including Random Forest, XGBoost, CatBoost, and Light Gradient Boosting Machine (LGBM). These are well-suited for genomic data as they can handle high-dimensional, discrete, and missing data in an accurate and fast manner. Furthermore, their non-linearities can capture dependency information generated by linkage disequilibrium (LD) within the sequences.
- Linear Models: including Logistic Regression, Linear Discriminant Analysis, and Naive Bayes. Linear models provide fast and interpretable classifiers that can make use of the divergent SNP frequencies between populations to successfully classify each window.
- Support Vector Machines and k-Nearest Neighbour: including SVMs with linear kernel, radial basis function kernel, string kernel, and our novel CovRSK. Kernel Machines such as SVMs and k-NN can accurately classify genomic sequences by computing a similarity measure between the input sequence and some support (training) sequences of known ancestry. Furthermore, the string kernel and CovRSK can handle admixture and linkage disequilibrium (LD).

#### 2.1.3 Smoother module

After the windowed SNPs are passed through the base module, the initial ancestry estimates for each window are refined with a smoothing module Figure 1e-h. We introduce a completely data-driven approach to produce refined ancestry estimates using discriminative smoother models that are trained to fix the errors made by the base module classifiers and that learn the statistical properties of ancestry switches (recombination probabilities) along the chromosome without needing information about the dates of admixture events (prior on ancestry segment lengths) or other model based assumptions. Such demographic parameters are typically not known exactly and may also vary across samples in a cohort. This approach differs from previous methods like RFMix^12^ that do require explicit specification of time-since-admixture and employ the same constant to represent the transition probability between all population pairs. Our trained smoother allows transition probabilities to be inferred (implicitly or explicitly - depending on the smoother) for each ancestry combination and allows these probabilities to vary for each window of the genome, resulting in far more flexible modelling and obviating the need for specifying a single (generally unknown) time-since-admixture parameter. Except for the CRF, which lacks it, our smoothers share only one hyperparameter, the smoothing window size, which specifies how many surrounding base classifier windows to include as input to predict the final ancestry estimate for a given window.

However, simply passing available training data through the base models, acquiring the probability estimates, and then using them as input in the learning procedure for the smoothing module, introduces a subtle bias; namely, the distributional shift at test time:

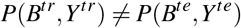

That is, the joint distribution of the base module probabilities, *B,* and the correct labels on the training data, *Y*, will in general be different during training and inference (see Figures A1 and A2). This is because the base module will be over-fit to the training samples and thus produce overconfident predictions. To avoid this, we split the available training data into two sets: one used to train the base module and one to train the smoothing module. The smoother training sequences are first passed through the base module to obtain unbiased (not over-fit) estimates of the probabilities that, along with the corresponding labels, better resemble the joint distribution at test time. These unbiased probabilities are used as training input for the smoother module. Note that this reduces the amount of training data available for both modules. However, training the smoothing module proves to be data efficient, since it can learn across the entire genome, and experiments show that its performance is not affected by the training data reduction (see Figure A3). To avoid a degradation in accuracy of the base module, we re-train the classifiers on the complete training data once the smoothing module has been fully trained. Supporting analysis can be found in Appendix B and the full pipeline is elaborated in the Methods section.

We explore three different types of data-driven smoothers:

- Linear Convolutional Filter (Logistic): a linear convolution followed by a softmax activation (a convolutional multinomial logistic regression) trained with cross-entropy loss learns a smoothing filter that combines probabilities predicted within neighbouring windows to obtain more accurate final probabilities. Linearity provides interpretable predictions and the weights of the smoother can provide insight into the structure of the training data (see section 2.4).
- XGBoost: boosting is well suited for the task of aggregating and refining probabilities. XGBoost is applied in a sliding window (convolutional) fashion with the same model applied to each set of windows. Since XGBoost is nonlinear, it can capture more complex dependencies between windows, leading to more accurate predictions.
- Linear-Chain Conditional Random Field: CRFs were used in RFMix, so we revisit them as smoother functions. Here we learn the transition probabilities by gradient descent instead of using priors.

#### 2.1.4 Calibration

Gnomix outputs ancestry probability estimates for each window. By selecting the ancestry with highest probability, an ancestry label can be assigned at each window. In order to interpret these probability estimates as confidence measures we need the system to be calibrated, so that the predicted probability approximates the true probability for each ancestry label. In other words, the predictions given a probability *X*% are correct *X*% of the time. However, most classifiers do not satisfy this property; thus, we include a calibration step that transforms the predicted probabilities (smoother module output) into calibrated probability predictions. Specifically, we use isotonic regression^21^, a non-parametric method that assumes the probabilities are a monotonic transformation of the predictions. By using isotonic regression, the probabilities are shifted closer to the true probability for each ancestry label. The calibration module is trained with the base model training subset (data not used by the smoother during training) and evaluated on the test dataset. For quantitative evaluation metrics, see Appendix C and Figure A4.

### 2.2 String kernels

In 2000 Lodhi et al.^20^ introduced the string kernel, an inner product in the feature space that is generated by all subsequences of a certain length. For two sequences of length *M*, 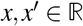, the string kernel is given by the canoncial equation:

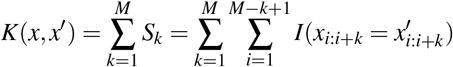

where *S_k_* is the number of subsequences of length *k* where *x* and *x′* are equal, or in other words the total count of k-mers that match between the two input sequences. This kernel has proven especially suitable for genome sequence comparison including for LAI^22, 23^.

#### 2.2.1 String kernel regularization

Despite its advantages, the string kernel has a few computational inefficiencies and weaknesses. For instance, it can be shown, by considering the subsequence equality counts *S_k_* as features that generate the kernel feature space, that the string kernel features are highly correlated given certain assumption. In particular, by denoting *q* as the probability that an input feature is the same for two individuals, one can model the kernel features as binomials,

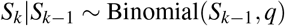

The covariance between two kernel features is then given by,

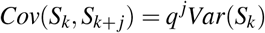

which can be used to bound the correlation from below,

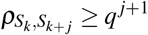

See sections E.1, E.2 and E.3 for more details. In the case of genomic sequences, q is often at the high end of the [0,1] interval, resulting in a high correlation between close kernel features.

The variances of the string kernel features decrease dramatically with the subsequence length *k* (see Figure A5) and as a result, their contribution to the feature space decreases and redundancy increases. That is, larger subsequences are less likely to affect the similarity measure of two sequences when included. Given the high correlation between neighbouring string kernel features, the effectiveness of a given feature is also highly dependent on its neighbouring features being included in the kernel (see Figure A6). We use these two observations to produce a more accurate version of the string kernel for this application, which we call covariance reducing string kernel (CovRSK).

CovRSK uses dynamically weighted sampling to select the features to include in order to reduce redundancy and feature correlation. The sampling procedure starts with an empty set of features and, in increasing order, adds a feature k with probability:

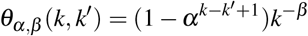

Here, *k^−β^* corresponds to the decaying property of the feature variance and the first term, (1 – *α^k−k′+1^*), is a discount proportional to the lower bound on the correlation with the last feature added and the one being considered, 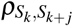. Both of those terms serve the same purpose; making more redundant features less likely. The hyperparameters *α, β* ∈ [0,1] affect the strength of the covariance reduction and the number of features sampled, respectively. It should be noted that CovRSK can be seen as a generalization, since with *α* = *β* = 0 it recovers the original string kernel. The technique is given in more detail in Algorithm 1.

The inclusion of string kernel features that are spatially correlated does not result only in redundancy, but also leads to an overfitting to longer matches leaving the model sensitive to sequencing errors (noise) that can break up the longer matches. In sections E.4 and E.5 we demonstrate how CovRSK acts as a regularizer for the string kernel resulting in robustness to such noise from genotyping call errors (Figures A7 and A8), and we visualize an example of the sampling procedure in Figure A9.

##### Algorithm 1: CovRSK

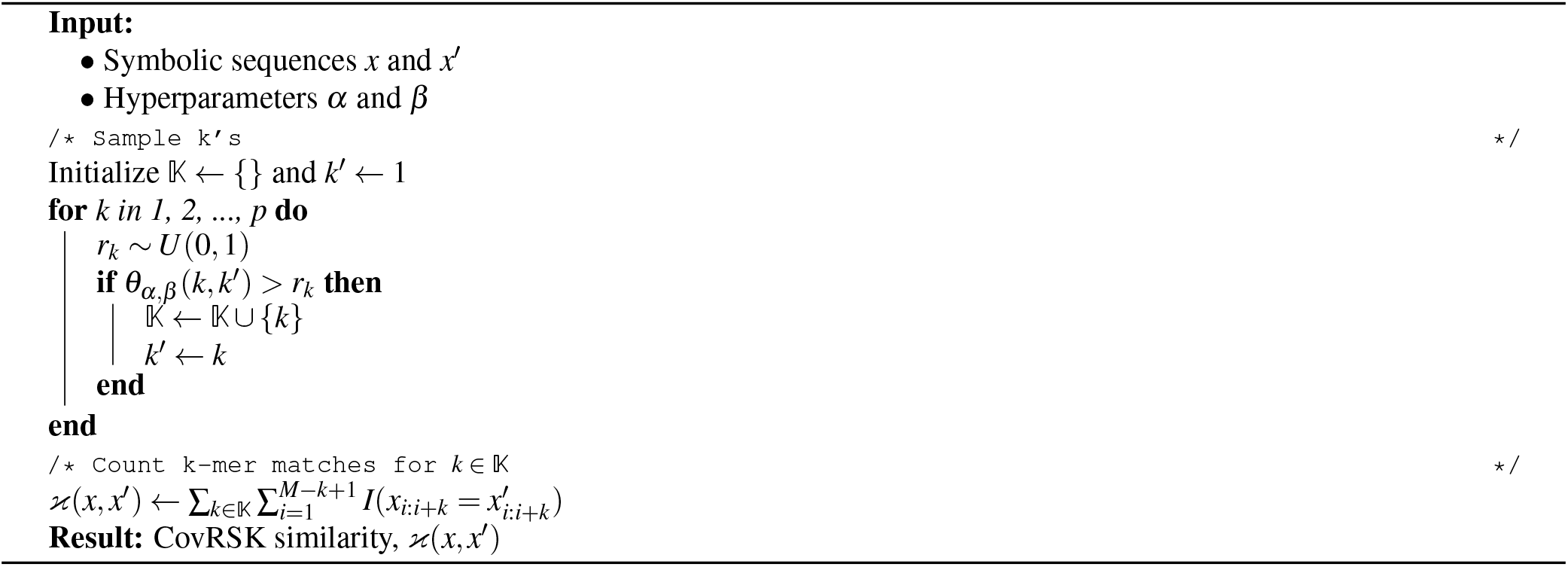

#### 2.2.2 Efficient computation of the string kernel

From the canonical string kernel equation above in section 2.2, containing an inner product between two sequence vectors, it is evident that a naive computation would grow quadratically with sequence size. However, unless these two inputs are a perfect match, one does not have to check all of their subsequences. If a subsequence of length *k*, starting at certain position, is not a match, then all sequences of length *k′* > *k* starting from that same position will also not match. Thus, a dynamic programming (DP) algorithm, which scans for matches only when shorter matches have already been confirmed, is appropriate. This lends itself to recursion, a longer match-check subroutine is called only after finding all shorter matches starting from same positions. However, without caution, this can result in an algorithm that still has quadradic complexity, which is easy to see in the worst case when the input sequences match exactly. Here one needs 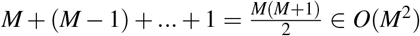 additions, one for each matching k-mer.

A better approach follows from the realization that each contiguous matching subsequence *μ* of the two sequences contributes independently to the total similarity,

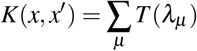

where *λ_μ_* is the length of match *μ* and *T*(*λ*) is the total number of k-mer matches within a contiguous match of length *λ*. Now the number of k-mer matches within a given match of length *λ* is the triangular number of that match length *T*(*λ*), denoted as *T_λ_*, and computed by

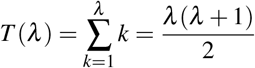

This inspires a simple, linear time algorithm for computing the string kernel that is shown in Algorithm 2 and illustrated in Figure 2.

##### Algorithm 2: String kernel with triangular numbers pseudo code

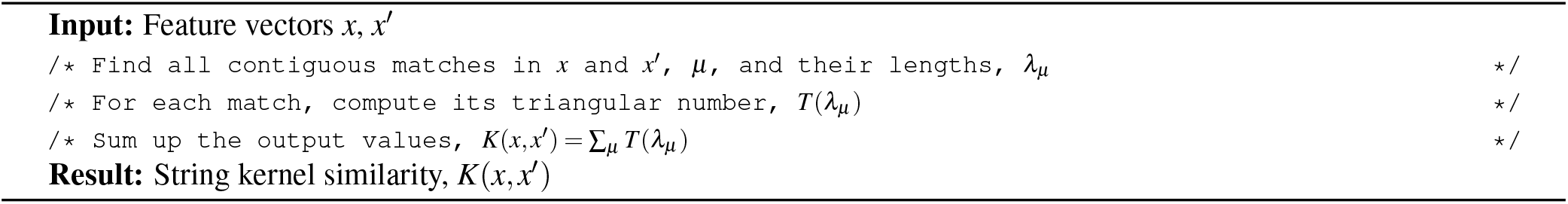

**Figure 2.**
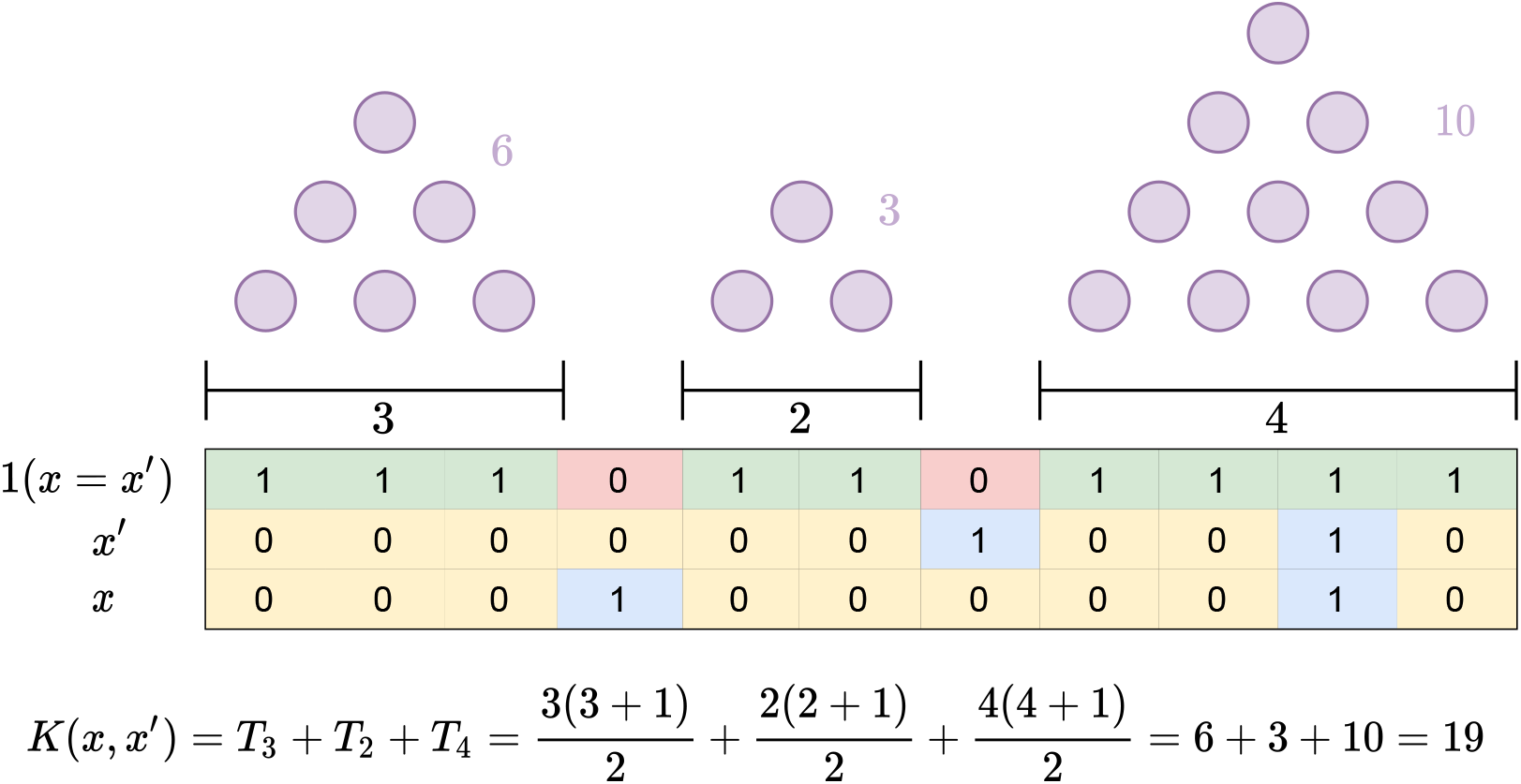
String kernel computation with triangular numbers visualized for two sequences, *x* and *x′*, of length eleven.

Computing the kernel for a data matrix of *n* individuals requires *n*^2^ such kernel similarity scores. For further computational optimization, we implemented a DP version of Algorithm 2 that can be vectorized across samples. This version is adjustable for efficient computation of the CovRSK. Detailed implementations of the string kernel with triangular numbers, with the DP version, and with the CovRSK customization are listed in Algorithms 5, 6 and 7.

#### 2.2.3 String kernel generalization and kernel comparison

The string kernel computation can be considered as a summation of polynomials *K*(*x, x′*) = ∑_*μ*_ *g*(*λ*): we find all contiguous matches of the input sequences and their lengths *λ*, compute the polynomial 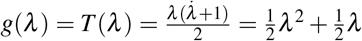, and sum the outputs. The question arises as to whether this polynomial is particularly suitable, or are better functions for the task. For instance, the Hamming kernel (another kernel that is historically popular within bioinformatics^24^) given by 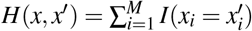 could be computed in the same manner using the trivial linear polynomial, *g*(*λ*) = *h*(*λ*) = *λ*. Intuitively, the choice of polynomial affects the contribution of longer matches relative to shorter ones. In the case of the Hamming kernel, longer sequences have no more importance than shorter ones, while the canonical string kernel weighs sequence length contributions quadratically.

This question leads to a generalization of the string kernel which replaces the triangular number polynomial, *T*(*λ*) with other polynomials *g*(*λ*). We refer to this generalization as the Polynomial String Kernel and describe it in Algorithm 3.

##### Algorithm 3: Polynomial string kernel pseudo code

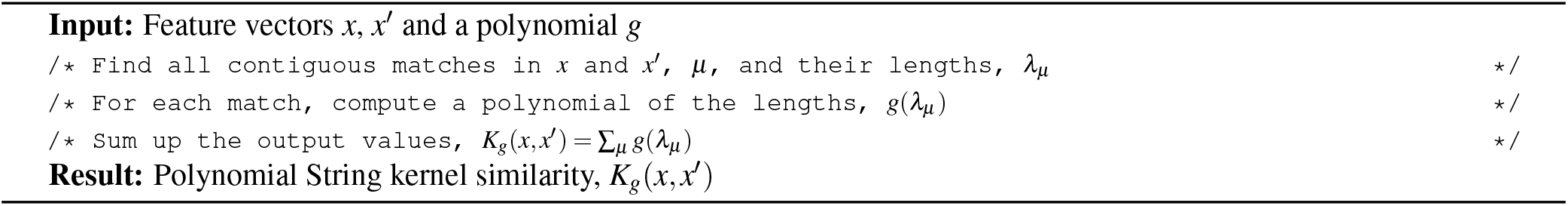

We investigate the optimality of the String Kernel empirically in Figure 3. We plot the validation accuracy for an SVM with different instances of the Polynomial String Kernel consisting of monomials of degree between one and two and note that neither the Hamming kernel, *g*(*λ*) = *λ* nor the string kernel, 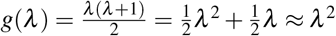, is optimal. Instead we see the optimal degree for a monomial instance of the Polynomial String Kernel is approximately *g*(*λ*) = *λ*^1.2^, nearly reaching the accuracy of CovRSK.

**Figure 3.**
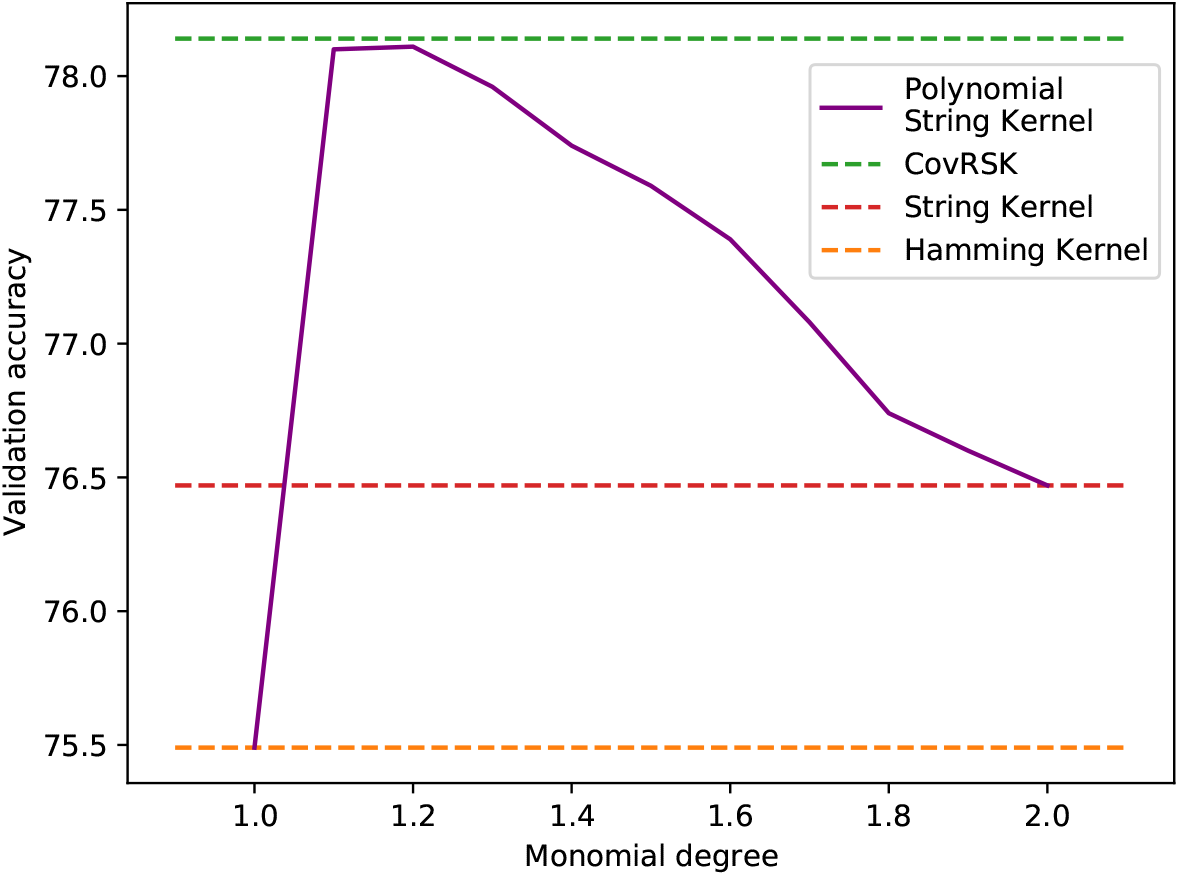
Validation accuracies for SVMs with different kernels on the dev dataset. The optimal degree of monomials is between 1 and 2, suggesting that neither the Hamming kernel (monomial of degree 1) nor the String kernel (polynomial of degree 2) are optimal. CovRSK has the highest accuracy, closely followed by the optimal monomial instance of the Polynomial String Kernel.

### 2.3 Experimental results

For thoroughness and a fair comparison to existing methods, we generated multiple continental-admixed datasets. We report accuracy, training time, inference time, and model size for our various model configurations on these datasets, which we refer to as the Latin American, 5-ancestry and 7-ancestry sets. See Table 1 for a detailed breakdown. The data used for each is whole genome, but for completeness we also subsampled a fraction of the SNPs corresponding to those positions found on a common genotyping array (specifically that used in the UK Biobank) to create another benchmark dataset (array genotyped). Thoughout we consider chromosome 20. We compare all of our results with the most widely used current LAI methods, RFMix (Random Forests)^12^, ELAI (HMMs)^13^, and Loter (dynamic programming)^14^, showing that our method widely outperforms each in both accuracy and speed on whole genome data. These results are presented below and in Appendix D.

**Table 1.**
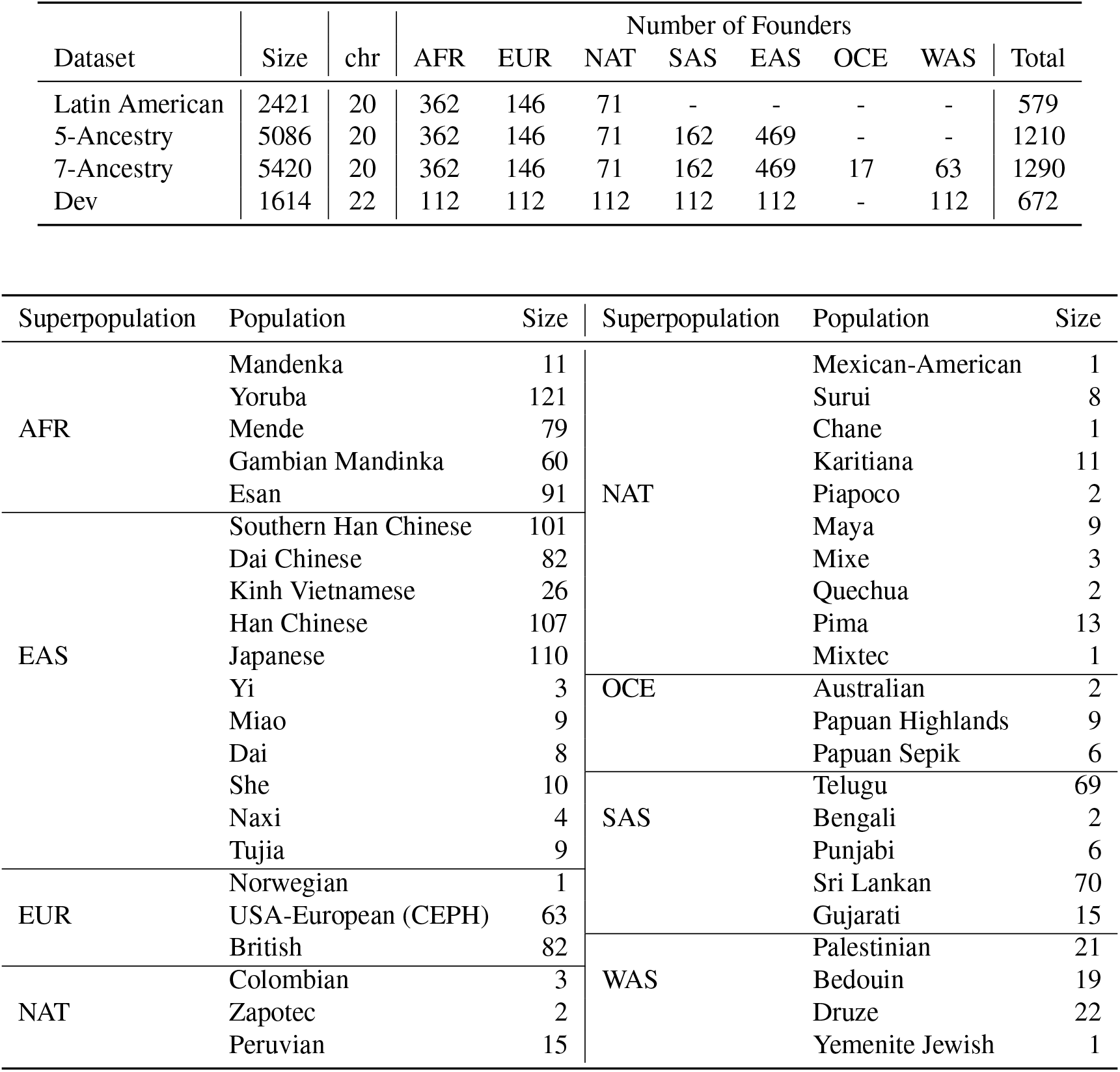
Dataset compositions.

First, we performed an evaluation of multiple classifiers of the base module. Specifically, we evaluate random forest (RF), CatBoost (CB), XGBoost (XGB), light gradient boosting machine (LGBM), K-nearest neighbors (KNN), support vector machine (SVM) with radial basis function, SVM with a string kernel, SVM with the polynomial string kernel, SVM with CovRSK, linear discriminant analysis (LDA), logistic regresssion, Multinomial Naive Bayes (MNB), Binomial Naive Bayes (BNB) and Gaussian Naive Bayes (GNB). Figure 4c compares these accuracies and training and inference times. We see that CovRSK SVM is the most accurate classifier, although with a large training time and inference time. Following CovRSK, polynomial string kernel and logistic regression provide comparable accuracies with the latter the fastest. Indeed logistic regression has a competitive accuracy, surpassing the other linear models including LDA and Naive Bayes, and the fastest running time of all models. We note that the gap between Naive Bayes, which assume independence between features, and logistic regression, which does not, indicates the importance of accounting for the correlation (linkage) between SNPs. Similarly, the additional accuracy of string kernels, which can learn linkage dependencies, over linear models, shows the importance of these dependencies, particularly for lower resolution array genotyped datasets (4c). Tree-based methods are seen to have a wide range of accuracies and training/inference times. RF provides much lower accuracy than the boosting-based methods, while XGBoost provides the most accurate predictions among the tree-based methods with the trade-off of the largest training and inference time. Overall, tree-based methods perform poorly compared to kernel and linear methods. Furthermore, linear models are considerably faster than tree-based models and kernel-based models. Given these results we select two classifiers with different accuracy/speed trade-offs: SVM with CovRSK as the most accurate - but slow - of our classifiers, and logistic regression as an accurate, interpretable, and fast classifier.

**Figure 4.**
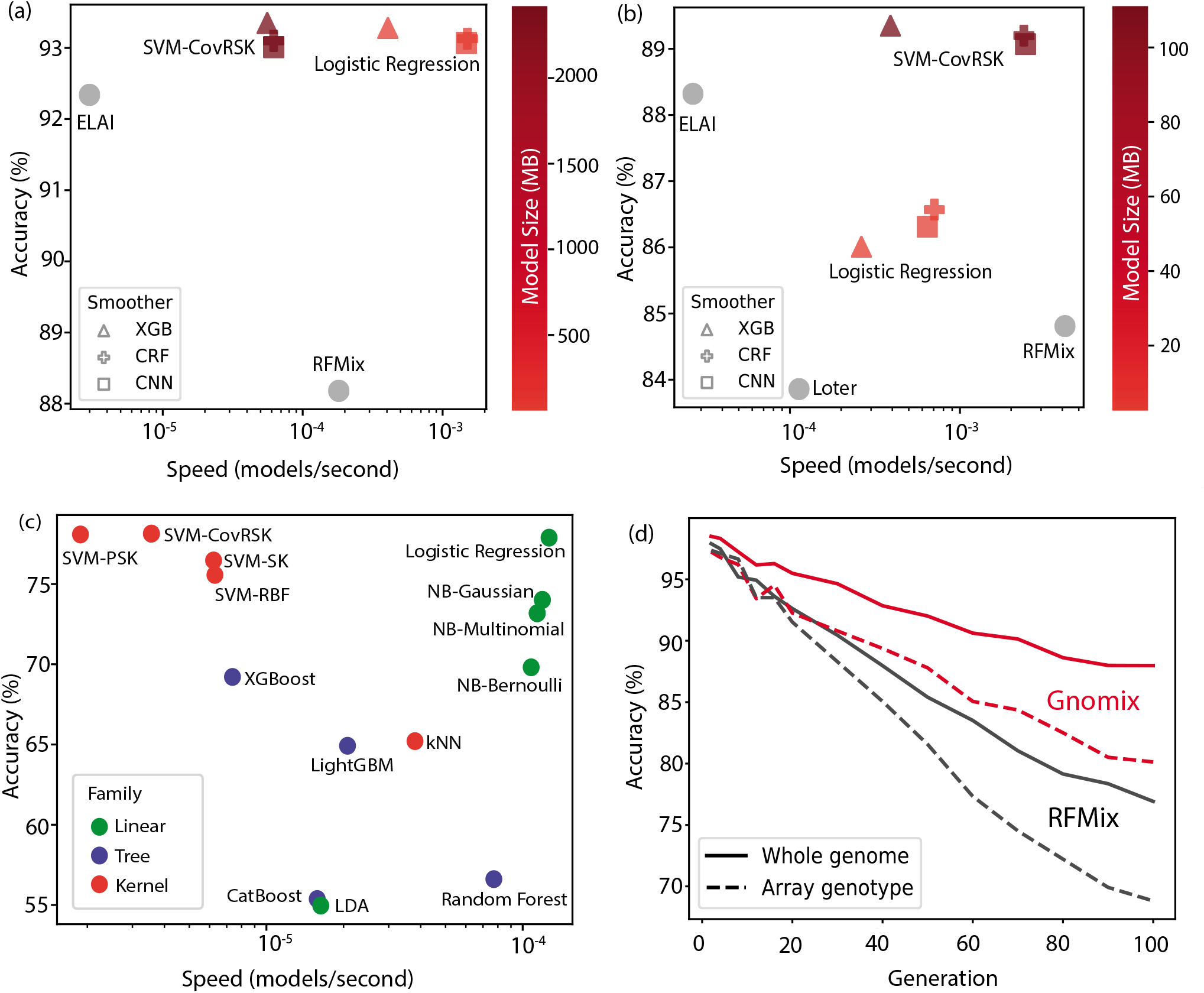
Performance plots. **(a)** Performance on whole genome data. Accuracy, training time and inference time of our models (size of model indicated by shade of red) and existing algorithms (shaded grey, since they do not permit saving and sharing of models) on the seven ancestry whole genome data, showing our models beating existing algorithms by a wide margin. Loter could not run on whole genome due to excessive memory requirements. **(b)** Performance on array genotyped data. Same analysis as (a) for models trained using only the SNPs found on the UK Biobank Axiom genotyping array. **(c)** Comparison of our different base models on whole genome data. A pareto optimal frontier is described by our models, SVM-CovRSK and Logistic Regression. **(d)** Accuracy into the past. Accuracy of our best configuration, CovRSK SVM with XGBoost smoother (solid line), versus RFMix, the existing method with practical speed on whole genome (a), as a function of number of generations since admixture. Larger numbers of generations since admixture generate more ancestry switches and smaller average lengths of ancestry segments, leading to a more challenging problem. We see that Gnomix has higher accuracy than RFMix even when Gnomix has access only to low resolution data (array genotyped, dotted line), and RFMix has access to full resolution data (whole genome, solid line).

We perform an evaluation of different methods used as smoothing modules using SVM with CovRSK and logistic regression as base module classifiers. Figure 4a (whole genome) and Figure 4b (genotyped array data) show the training and inference speed and the accuracy of the different combinations of classifiers and smoothers. We see that CRF and convolutional smoothers are consistently faster than XGBoost smoothers, but typically with a lower accuracy. Additionally, while CovRSK is slower than logistic regression on whole genome data, it provides improved speed when working with genotyped array data. We additionally compare these different configurations of our system with the currently most common LAI methods: RFMix, Loter and ELAI. We observe that our CovRSK-XGboost surpasses all of these methods in accuracy. Indeed, all of our configurations surpass RFMix in accuracy, and almost all surpass Loter and ELAI in training and inference time (on both whole genome and on genotyped array data). On whole genome data all of our configurations surpass existing methods in speed and in accuracy by a large margin. In particulary, our CovRSK-XGboost combination outperforms all others in accuracy, albeit with the downside of larger training and inference time, while logistic regression, a close second in accuracy on whole genome, provides a very fast alternative for processing massive datasets.

We note that the CovRSK provides a significant improvement with respect to logistic regression in UK Biobank genotyping array data, however, in whole genome both methods perform similarly. This seems to indicate that with enough SNPs-per-window, in other words, enough input features for each base classifier, linear models such as logistic regression perform extremely well with more computationally complex non-linear techniques, such as CovRSK, providing only marginal improvement. However, in scenarios with a low number of SNPs per window, non-linear models that generate a richer feature space, such as CovRSK, provide a significant improvement on accuracy.

Robustness to the generation time since admixture is another important aspect of LAI methods. This generation time will determine the length distribution of the segments originating from each ancestry group, so it is important that methods can properly deal with a wide range of generation times and length distributions. We compare our top performing configuration (CovRSK SVM with XGBoost smoother) with RFMix (the only existing method with practical run-time on whole genome Figure 4a) over a wide range of generation times Figure 4d. We can see that RFMix accuracy decreases rapidly as the generation number increases (segment sizes decrease), while Gnomix maintains a good classification accuracy (>90%) well beyond 100 generations (approximately 3000 years ago for humans).

In addition to being able to handle multiple generation time regimes, Gnomix is also robust to the population set that it is trained on. While some closely related populations, such as West Asian and European, pose a significant challenge for methods like RFMix, Gnomix handles these far better because its base models and non-linear data-driven smoothers are more accurate.

### 2.4 Population structure is reflected in the learnt models

The effect of the smoother for an example haplotype is shown in Figure 5a. Here the model makes errors on two segments that both - in their own way - shed light on the typical errors that can occur. On the left the model has difficulty determining the switch points (boundary between two ancestries). Indeed, by looking at Figure A10 it is clear that the model accuracy increases rapidly with distance from a switch points with most errors occurring at or around these points. The error on the right is a West Asian segment wrongly classified as European. This phenomenon will be seen again in the discussion of Figure 5b below, where the similarity of these two related populations plays a role in confusing the classifier. Indeed, given these two ancestries’ shared historical introgression from early farmers in the Middle East, this European prediction might stem from genuine shared ancestry. When observed closely, the independent base probabilities seem to be around the same magnitude for both West Asian and European in this segment, further supporting that hypothesis.

**Figure 5.**
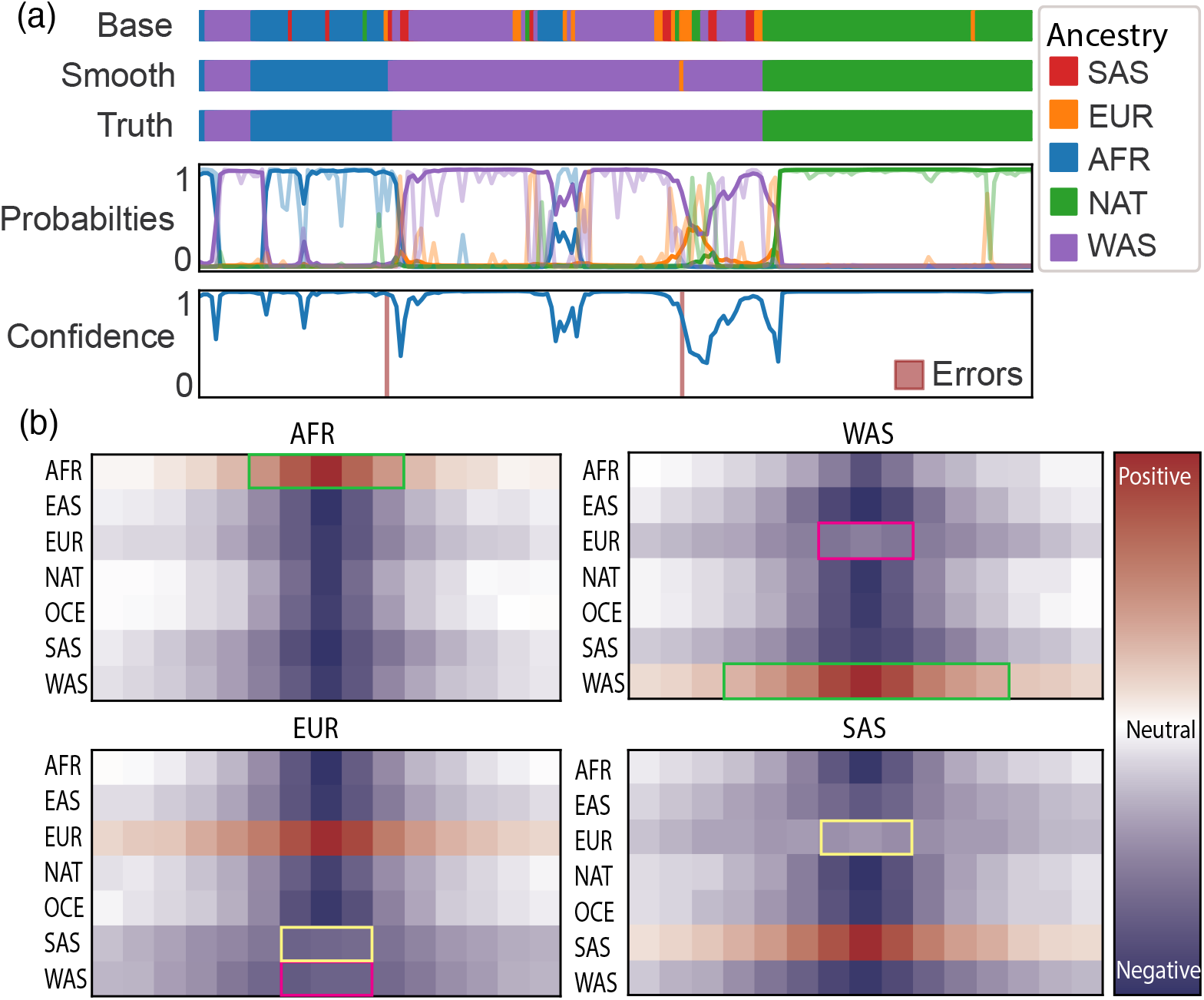
Effect of population structure on learnt models. **(a)** Analysis of predicted probabilities and errors of Gnomix on a sample admixed haplotype. The color for each ancestry is the same for the smoother and base probabilities, but the latter is faded. The model makes two errors; one at an ancestry switch point and one where it mistakes a small West Asian (WAS) segment for a European (EUR) one. **(b)** Weights (importance measure, red positive and blue negative) for the base model predictions for each ancestry (vertical axis) for each window (horizontal axis) learned by the convolutional smoother to predict each ancestry (plot titles). Model trained on seven populations: African (AFR), East Asian (EAS), European (EUR), indigenous American (NAT), Oceanian (OCE), South Asian (SAS), West Asian (WAS). Note the African smoother does not need to consider many surrounding African window probabilities (green rectangle, upper left), because this ancestry is so well separated from the others, while the smoother for West Asian ancestry, which is not well separated from European, needs to consider more surrounding windows (green rectangle, upper right). Shared introgression (perhaps Early Farmer^25^) between West Asian and European ancestries results in only weakly negative weights for European probabilities for the West Asian smoother (pink box, upper right), which must then be aggregated across many more surrounding windows, and similarly for European and South Asian (yellow box, lower right) due in this latter case perhaps to shared Indo-European introgression^26^.

Analyzing the weights learnt by the linear convolutional model, which indicate how the smoother module aggregates information from the base models’ outputs, we gain insight in Figure 5b into the data driven smoother. By learning a linear filter in a data-driven way, the magnitude of filter weights can be interpreted as a feature importance indicators. In Figure 5b, we can see from the linear smoothers’ filter weights that, in order to predict a particular ancestry in a window, the base models’ priors are taken strongly into account. There is a strong positive correlation between base model probability estimates and the smoother predictions of a given ancestry (weights colored red). Further, one can see that the spread of positively correlated windows is higher for populations that are less well separated, like West Asian (WAS) and European (EUR), as compared to population labels that are well separated, like African (AFR). Examining the center columns of the European (EUR) filter, we note that having a high base probability for EUR correlates positively with being predicted European (red), but is negatively correlated (dark blue) with being predicted AFR, OCE (Oceanian), EAS (East Asian), NAT (indigenous American). Interestingly, EUR is less negatively correlated (lighter blue) with predicting West Asian (WAS) and South Asian (SAS) than with any of the other ancestries, likely because both of these populations have shared historical introgression events with Europeans, namely the spread of early Near Eastern farmers in the case of WAS^25^, and the spread of the Indo-European languages in the case of SAS^26^. Furthermore, this effect is not observed between South and West Asians in our dataset, suggesting that the common ancestor between the European and West Asian references, namely neolithic Anatolian farmers, is different than the common ancestor between the South Asian and European references, namely Indo-European speakers.

The presence of such shared population histories adds complexity to this inference task and can help explain the success of nonlinear smoothers. We also note that choosing reference samples and label names in this context is not always simple and depends upon the time period being investigated. Although some continental populations have been long separated, diverging through independent genetic drift^27^, no human population is an “island” without historical introgression, admixture, and shared ancestry with others^28^. Care must thus be exercised when choosing label names for genetic clusters to avoid conflating them with socially constructed ethnicities^29^, which vary rapidly in time and across nations and can be entwined with political considerations (nationalism, irredentism, and resource access).

### 2.5 Phasing error correction: Gnofix

Because most sequencing technologies in use are unable to assign an accurate parental chromosome source to each sequencing read in a diploid genome, several statistical algorithms have been developed to assign each variant to its correct parental haplotype. These techniques, known as phasing, take advantage of observed correlations between neighboring variants in databases of reference populations. Some of the most commonly used methods include Beagle^30^ and SHAPEIT^31^. While these tools are highly accurate locally – that is, the probability of having two neighbouring variants correctly phased is very high – they occasionally fail, producing swaps between the two parental sequences. Thus, the probability of having two distant variants correctly phased converges to random chance as their separation increases. Fortunately, such errors can often be fixed with LAI. Given the additional information present in ancestry calls along the genome, such sporadic switch errors become clear. The most commonly used phasing error correction method using LAI that can handle multiple ancestries is found in RFMix (version 1, but not version 2).

We propose a novel method, Gnofix, that employs a trained smoother from the Gnomix module to find the most likely phase assignment of an individual’s two haplotypes. This is done by using the confidence of the ancestry estimates for a given sequence as a proxy for the probability of that sequence being from the observed distribution of human haplotypes. Starting with a window-based ancestry probability estimate for an individual’s maternal and paternal haplotypes (phased using an existing algorithm), these probabilities are leveraged to iteratively swap segments of each haplotype until the most probable one is recovered. Such input probabilities can be obtained from many LAI models, including RFMix^12^, LAMP^10^, LAI-Net^15^ and, of course, Gnomix.

The algorithm starts from one side and during one iteration works its way to the other. At each step it defines its scope to be a set of local consecutive windows of the same length as the trained Gnomix smoother size. It then extracts the ancestry probabilities that lie in this scope and explores different local phasing alternatives by exploring potential combinations of swaps between the two parental sequences. This results in a collection of swap combinations. Each item in this collection is treated as a proposal for the true sequence. The collection need not contain the complete set of all possible swap combinations, however. Indeed, that would create redundancy, since neighboring scopes have a large overlap for swap proposals. In addition, for any sizable smoother size, such an exhaustive enumeration of all swap combinations would be intractable. Thus, the collection of swap combinations is restricted to include only swaps at - or closely around - the center, where the smoother is most receptive; this reduces the number of proposals dramatically. Once extracted, these proposals are passed into the smoother to obtain predictions for the center window and - more importantly - prediction confidences. The proposal with the highest confidence is then selected and the scope shifted along the haplotype for analysis of the next position. Once the full chromosomal sequence has been traversed, the scope is shifted back to the start and the next iteration begins. Iterations continue until convergence, or until a chose upper bound is reached. A general outline is shown in Algorithm 4 and the process is visualized in Figure 6.

**Figure 6.**
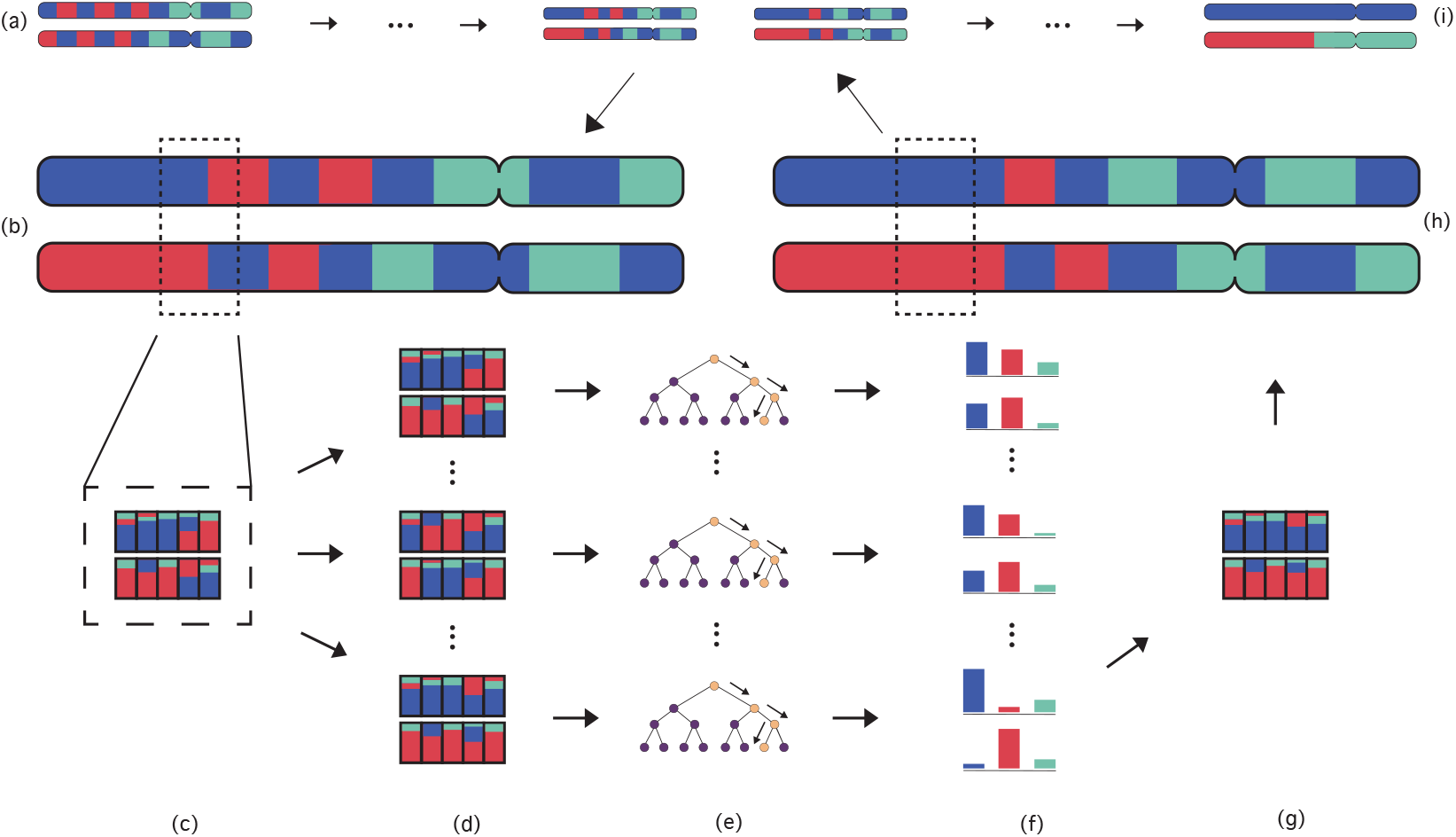
Phase correction algorithm. **(a)** Input consisting of maternal and paternal haplotypes containing phasing errors. The colors denote different ancestry predictions. **(b)** State of the haplotypes at a given state and then after some phasing errors have been corrected. Dashed box marks the scope with the center window. **(c)** Probabilities of each ancestry at each window in the scope are extracted. **(d)** Collections of swaps are proposed. **(e)-(f)** Proposals are passed through the smoother to obtain predictions and confidence. **(g)** Proposal with most confidence is chosen. **(h)** Windows in scope are returned back to the haplotypes. **(i)** Output maternal paternal haplotypes containing no phasing errors.

#### Algorithm 4: Gnofix

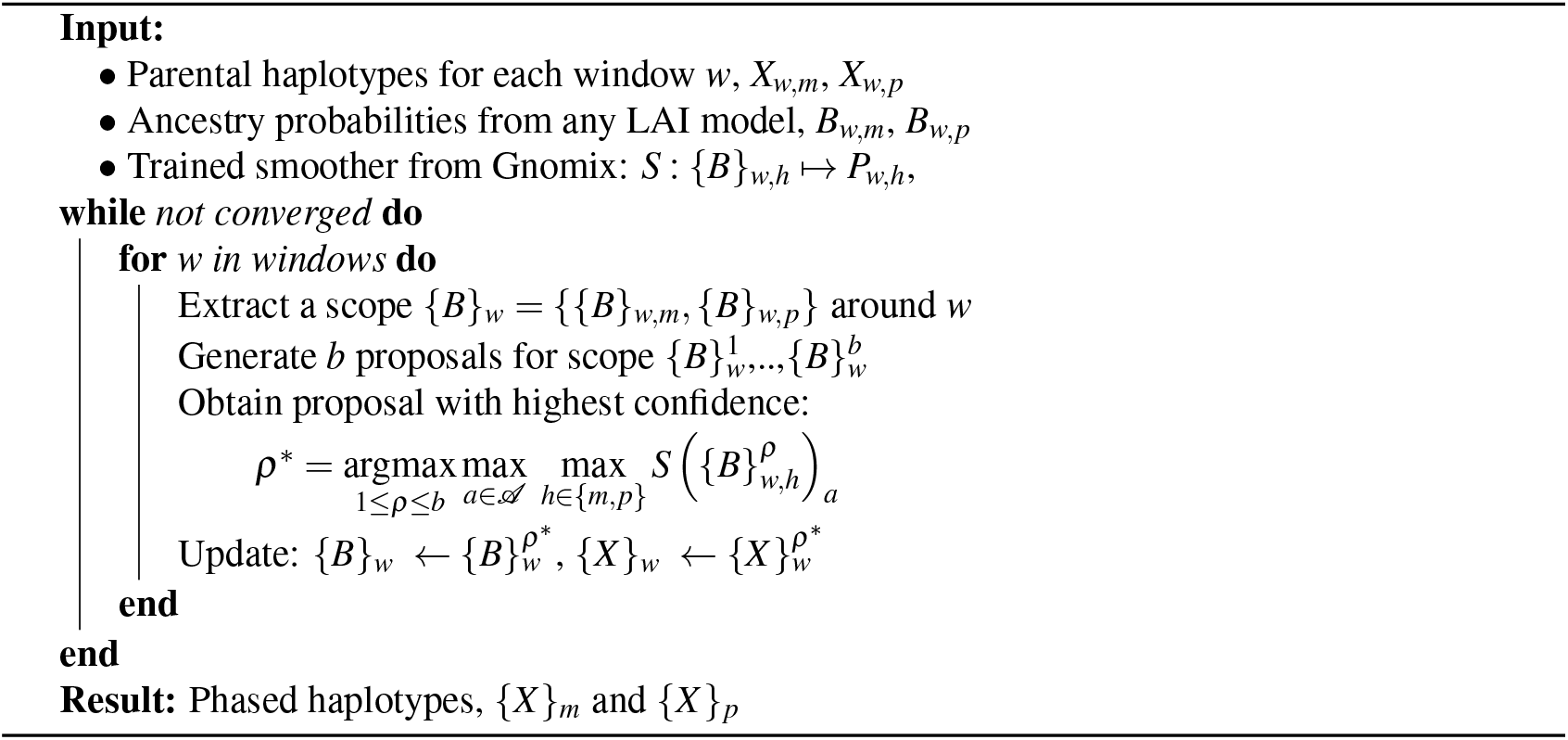

We compare the performance of Gnofix to Beagle phasing and to rephasing using RFMix by plotting the fraction of SNP-pairs correctly phased as a function of genetic distance between them, where the SNP-pairs were sampled randomly from heterozygous ancestry regions. (Homozygous ancestry regions cannot be phase-corrected using local ancestry.) When the data is not well phased we expect this fraction to decrease with genetic distance and converge to 0.5 (even chance). We perform this comparison on a Latin American phasing dataset created by simulating European, Native American, and African three-way admixture. RFMix and Gnofix algorithms show similar phase-correction accuracy (Figure 7); however, the run-time between them differs by several orders of magnitude. It takes Beagle and RFMix about 5 and 37 minutes, respectively, to run on this dataset; however, Gnofix takes simply 17 seconds. Such acceleration becomes crucial for large biobank datasets for which RFMix style phase correction becomes infeasible. See Appendix H for additional experiments.

**Figure 7.**
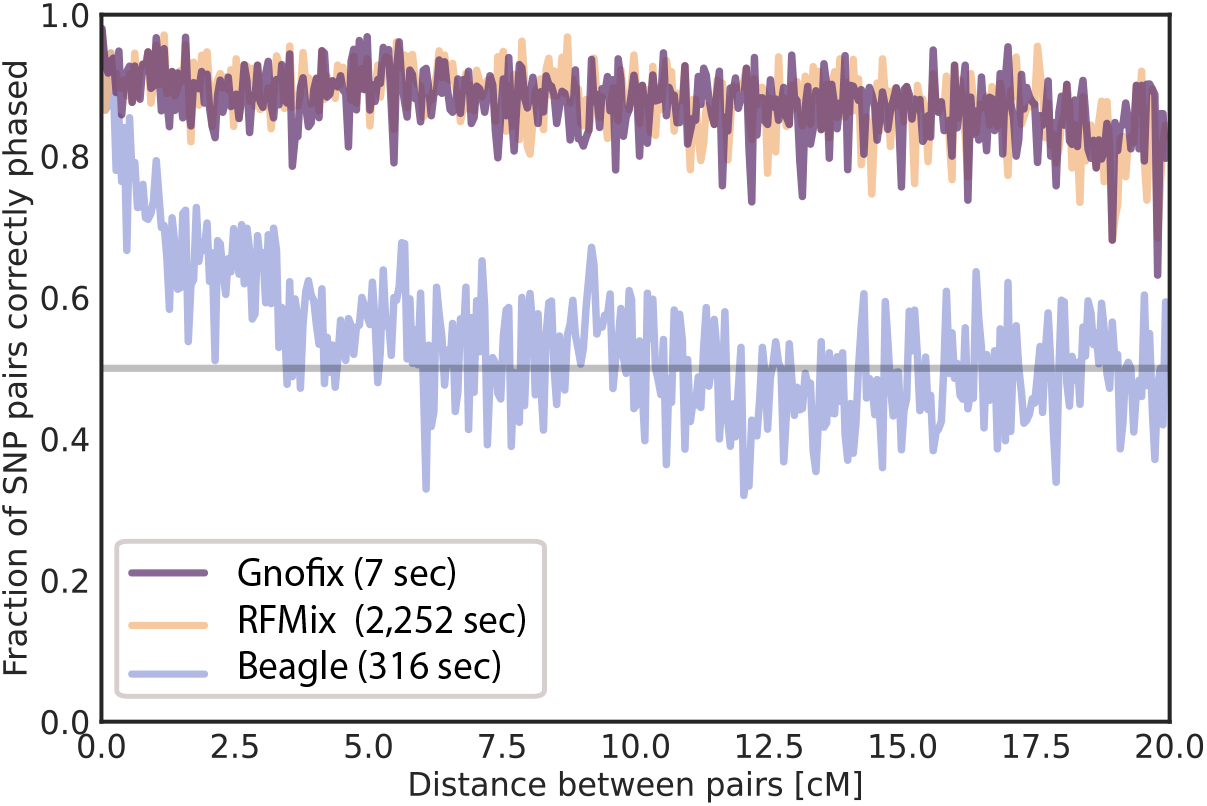
Re-phasing performance. Phasing error correction accuracy of Gnofix and RFMix on the Latin American phasing dataset.

### 2.6 Application to New World dogs

We use whole genome sequences of 191 canids from Plassais et al.^32^ to train a Gnomix model that can annotate canid species and breed along the genome. (See methods for more details.) We then demonstrate local ancestry calls on a panel of 531 dogs genomes obtained via the same study and finally use Gnofix to correct phasing errors.

We sort the local ancestry calls from each genome by total length of Arctic-breed segments called per sample and visualize the top ten dogs, see Figure 8a. We observe that all of these dog breeds, except for the Xoloitzcuintle (Xolo), belong to Arctic breeds or breeds known to be admixed with them (Spitz). Similarity in ancestry between Arctic breeds and Native American dogs has been noted previously; indeed, pre-contact Native American dogs lie closest to modern Arctic breeds in trees constructed from ancient genomes^33^. Thus, Arctic local ancestry models can also detect ancestry segments deriving from Native American dogs. Thus, these segments in the Mexican Xoloitzcuintle, which is said to have descended from pre-Columbian Mesoamerican dogs, may indeed originate in the pre-Columbian dogs kept by ancient Mesoamerican cultures (Figure 8b). The total length of Arctic ancestry segments that we identify in the Xolo amounts to 4.03% of its genome, which matches prior global ancestry approximations that estimated the Xolo’s pre-Columbian ancestry at 3%^34^. To our knowledge we are the first to identify this pre-Columbian ancestry at the genomic segment level.

**Figure 8.**
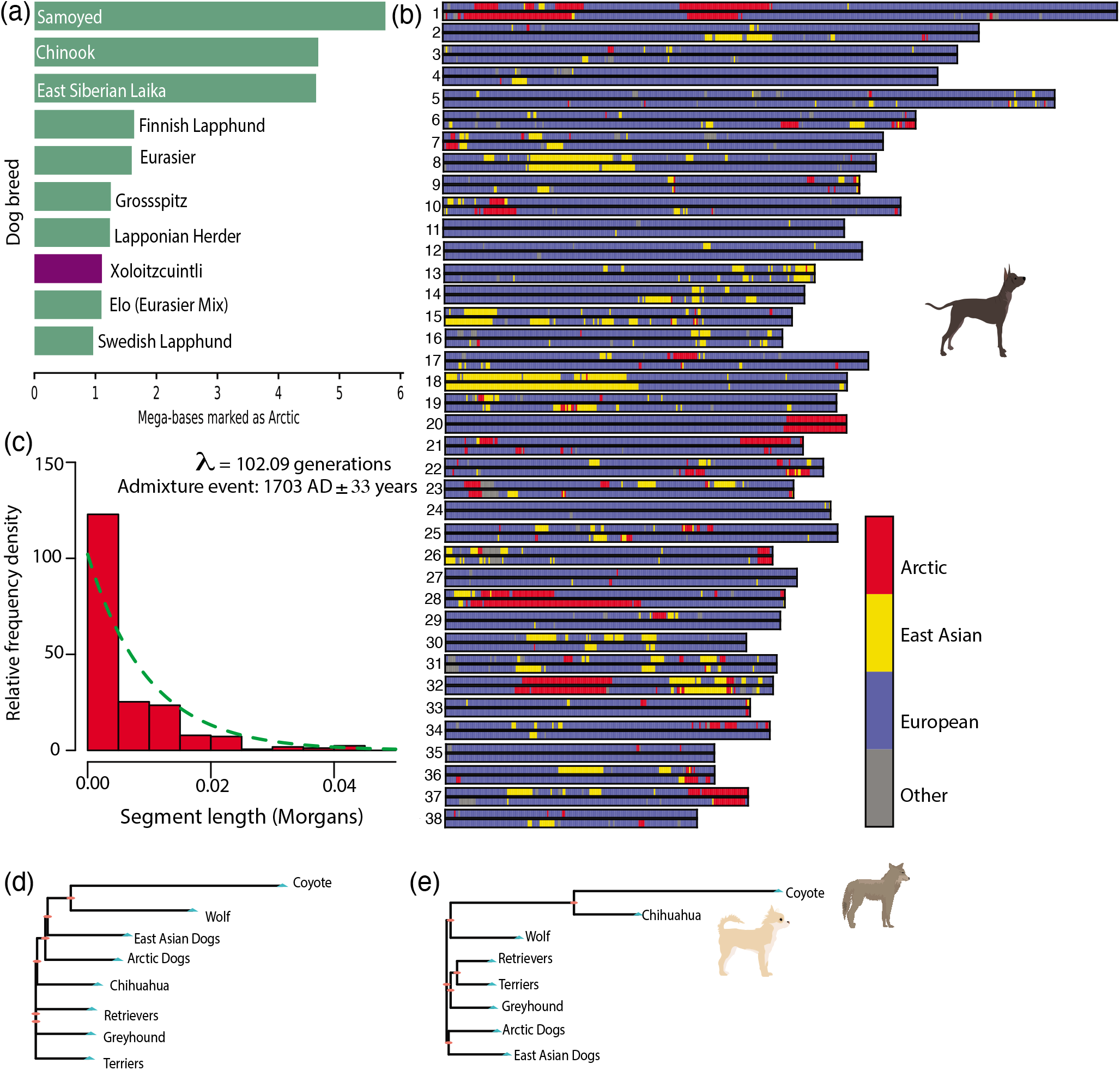
Gnomix applied to dogs reveals ancient New World admixture. **(a)** The ten breeds with highest total Arctic segment ancestry. **(b)** Chromosome painting of a Xoloitzcuintle (Xolo) dog. **(c)** Segment length distribution of Arctic segments in the Xolo. **(d)** Neighbor-joining tree for all of chromosome 1 for various canids using the average number of pairwise differences (*π*). **(e)** Neighbor-joining tree of *π* between coyote segments identified in the chihuahua by local ancestry inference versus sequences of other canids.

By fitting an exponential curve to the distribution of these Arctic genomic ancestry segments in the Xolo (Figure 8c) and using a generation time of 3 years for dogs^35^, we obtain an average admixture time of 306 years. This implies an average admixture date in Mexico between European and pre-Columbian dogs of 1704 AD ± 33 years, which notably matches the dating for admixture between European and pre-Columbian peoples in Mexico of 1746 AD (8.6 generations with 30 years/generation) as arrived at by Price et al. using a comparable single pulse admixture model^36^.

Another signature of New World dog ancestry comes from genetic introgression with the coyote^34^, a related canid (Figure 8d) found only in the Americas. There is evidence that pre-Columbian cultures cross-bred coyotes with dogs^33^, and we indeed find shared genomic segments between coyotes and the chihuahua, another breed with links to the Americas. In Figure 8e we display a neighboring joining tree for average number of pairwise differences (*π*) of sites called as “coyote” in a chihuahua’s chromosome compared against other dogs.

The capacity of Gnomix to identify and annotate short, ancient chromosomal segments in whole genomes with high resolution is unmatched by existing local ancestry algorithms (see Fig. 4d) and here allows us to unravel the history of New World ancestry in Mexican breeds of dogs at generation times (one hundred generations) that in humans would correspond to demographic events that occurred over 3000 years ago. Besides surpassing the capabilities of current LAI algorithms our method also takes only minutes, rather than days, to run on whole genomes.

## 3 Methods

### 3.1 Human datasets

We source our worldwide human whole genome data from three publicly available projects: 1000 Genomes^37^, Human Genome Diversity Project (HGDP)^34^, and Simons Genome Diversity Project (SGDP)^38^. We bring all data onto human genome build 37, retain only biallelic sites, phase using Beagle^30^, and then filter these samples to obtain 1,380 single-ancestry samples by running unsupervised ADMIXTURE clustering^39^. From the latter, eight well-supported ancestry clusters were selected by cross-validation: East Asian (EAS), European (EUR), indigenous American (NAT), Oceanian (OCE), South Asian (SAS), West Asian (WAS), Subsaharan African (AFR), and African Hunter and Gatherer (AHG). We retained only the first seven ancestry groups for our experiments, since AHG did not have enough training samples in our dataset for robust benchmarking.

For analyzing and evaluating LAI models, genetic sequences from real admixed individuals cannot be used, as the ancestry switch points along their chromosomes cannot be known without sequencing a pedigree (trios). We therefore simulate admixed individuals from sequenced single-ancestry individuals (founders) from various populations (Figure 9). Using these real individuals’ sequences, we simulate descendants of admixed ancestry using locus specific recombination rate parameters from the human genetic map^12^. In brief, following Karavani et al.^40^, the number of switch points is modeled as a Poisson random variable parameterized by the chromosome length in Morgans. Recombination positions are simulated uniformly along the chromosome using the genetic map. The simulation process can be seen as a “select-and-stitch” process, where a simulated individual inherits each segment of their chromosome from a subset of founders and the generation-since-admixture of the individuals determines the statistical properties of segment lengths. The select-and-stitch process allows for much more lightweight computation than complete generation-wise recursive simulation of the matching explicitly, while also avoiding the founder effects in admixed samples that result from such recursion with small population sizes per generation.

**Figure 9.**
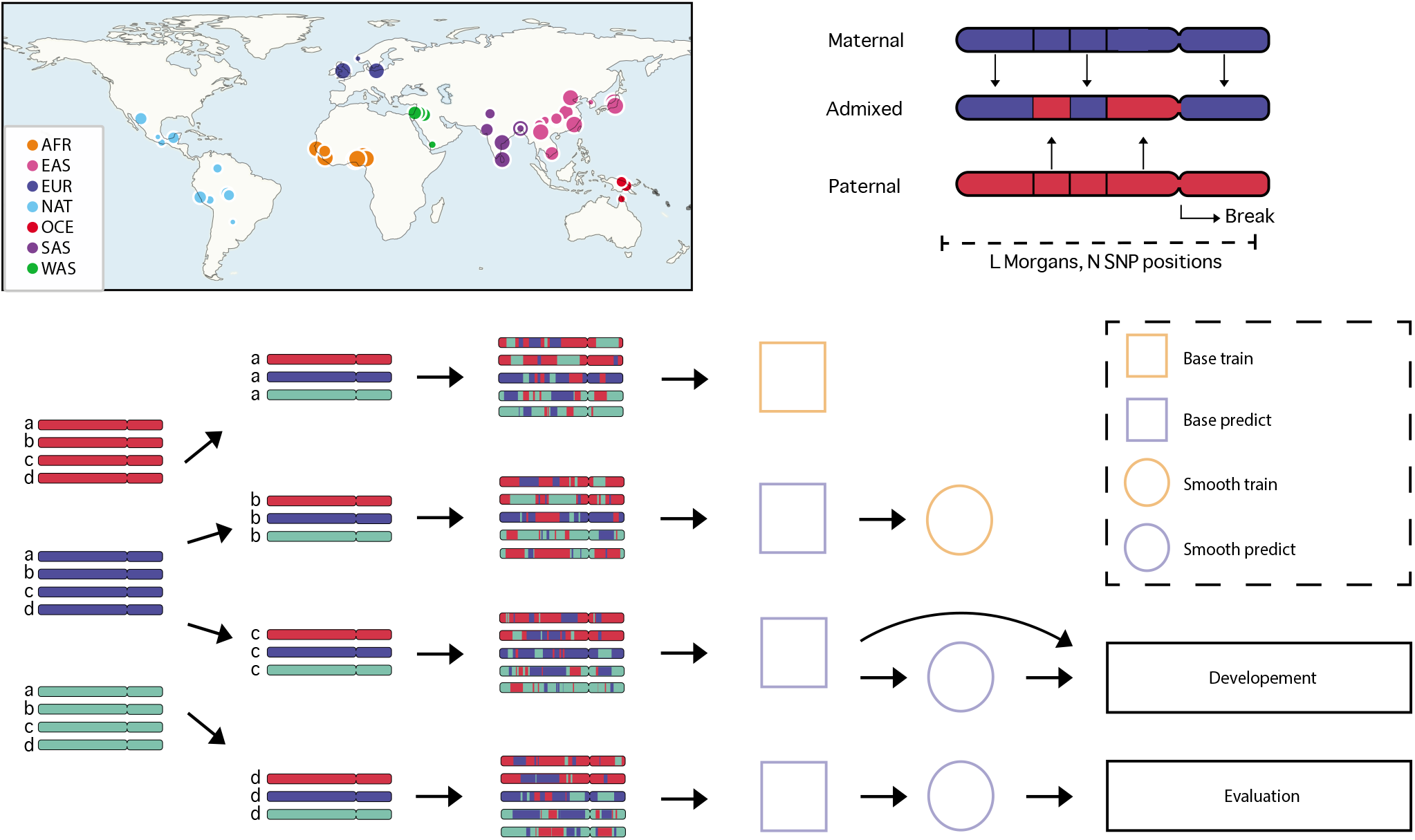
Origin and generation of human genome datasets. **Top left**, map of the distribution of single ancestry individuals used for dataset generation: African (AFR), East Asian (EAS), European (EUR), indigenous American (NAT), Oceanian (OCE), South Asian (SAS), West Asian (WAS). **Top Right**, simulated chromosomal crossovers between the maternal and paternal chromosome of a sample. The number of breakpoints are drawn from a Poisson distribution with a rate parameter dependent upon the generations-since-admixture, and break loci drawn uniformly along the chromosome. **Bottom**, dataset generation via simulation and splitting. The labeled single ancestry data is split between the independent sets and admixed individuals are simulated separately from each set to produce admixed individuals with labels. The base train data is used to train the base models and the smooth train data is used to independently train the smoothing module. Similarly the validation and test datasets are used to independently develop and finally test the model as a whole.

Gnomix development was performed on a development (dev) dataset: chromosome 22 of simulated admixed individuals from an even distribution of founders of six different populations, African, East Asian, European, indigenous American, South Asian and West Asian. 70% of the founders were used for training 12.5% validation and 17.5% testing. All splits contained admixed individuals of the generations 2, 4, 6, 8, 12, 16, 24, 32, 48, 64. On the dev dataset we compared various machine learning models for the base classifiers and smoother, where the hyper-parameters of the individual models were selected by performing a grid-search. We found values that worked well for the Gnomix specific parameters window size, smoother window size and context. While in theory these parameters could be application specific, in practice we find our models to be largely robust to these particular parameters on a wide variety of ancestries.

For the benchmarking datasets, we split the founders randomly in training founders (85%) and testing founders (15%). The Latin American dataset included founders with European (EUR), indigenous American (NAT), and African (AFR) ancestry. The 5-ancestry dataset had the founders from the same populations as the Latin American dataset, but added South Asia (SAS) and East Asia (EAS). While the 7-ancestry dataset added further Oceania (OCE, referred to as Papuan in some publications) and West Asian (WAS, anchored in the Middle East). See Table 1 for more detailed breakdown. Each model had access to the training founders and was evaluated on a simulated admix test set containing three times the number of samples as the testing founders. The generations used for testing were 2, 4, 8, 12, 16, 20, 30, 40, 50, 60, 70, 80, 90, 100 yielding 406 admixed diploid individuals equally split across the above generations.

Window size for Gnomix was one thousand SNPs for whole genome data and thirty SNPs for the array data, and a context of same size was used. The smoother size was 75 windows and the model output was calibrated with isotonic regression. The simulated admixed individuals used for training were of generations 0, 2, 4, 8, 16, 24, 32, 48, 64, 72, 100 and each population sampling probability was weighted to provide class balance. The default settings were used for RFMix, ELAI and Loter, and all methods had access to 32 CPUs.

### 3.2 Phasing Evaluation datasets

We simulate the phasing datasets of eighth generation admixed individuals using Algorithm 8 and then, after converting to genotypes, Beagle 5.1 is used for phasing. The Latin American phasing dataset was simulated from the validation founders with an even distribution of indigenous American, African and European ancestry: eight founders from each population. 67 founders from each population were used for training. Gnofix used a smoother that was trained on this dataset with a window size of 0.2 cM. It considered swap collections that had a single swap five or fewer windows away from the center. Lastly, we iterated through scopes where the base module probabilities were discontinuous. Since RFMix version 2 cannot perform phasing error correction, we used RFMix version 1. We used the default parameters with the window size also at 0.2 cM.

### 3.3 Canid Datasets

For training Gnomix on canid whole genomes, phased first using Beagle^30^, we used the following groupings as founders (sample numbers for each type in brackets): Coyote [3], Wolf (Grey Wolf [45] and Iberian Wolf [1]), Basenji [4], Tibetan Mastiff [10], Border Collie [15], Bull Terrier [9], German Shepherd [15], Greyhound [9], Hound (Afghan Hound [3] and Saluki [3]), Retriever (Labrador [24] and Golden [20]), Arctic (Alaskan Husky [2], Siberian Husky [4], Greenland Dog [1], and Alaskan Malamute [3]), East Asian (Shiba Inu [2], Chinese Shar-Pei [2], Jindo [1], New Guinea Singing Dog [4], Chongqing Dog [1], Xiasi Dog [1], and Chow Chow [4]). These groups were chosen based on the neighbor joining tree in Plassais et al. in order to ensure broad coverage from each clade of the tree^32^. With these groupings, we trained a Gnomix model for each of the 38 canid autosomes with the following configuration: logistic regression base, XGBoost smoother, 0.2 cM window size. We obtained an overall validation accuracy of 90.63%. This lower accuracy can be attributed to the only single digit numbers of whole genome training references available for many of the groupings (see above). Our >90% accuracy in the face of such paucity of references illustrates the ability of Gnomix to perform even when labelled training data is scarce, a case often encountered.

To produce the tree in Figure 8, we used neighbor joining^41^ with a pairwise distance matrix created by using the average number of pairwise differences (*π*) between various breeds as a dissimilarity measure^42^. The standard error for the number of generations since admixture was computed by performing one thousand bootstrap replicates and then refitting the exponential each time using maximum likelihood.

## 4 Data & code availability

The code for Gnomix is available at https://github.com/AI-sandbox/gnomix along with a user guide, documentation, pre-trained models, and directions to train on references of choice. The input datasets for our human genome modeling and analyses are available publicly through the Human Genome Diversity Project (HGDP), Simons Genome Diversity Project (SGDP), and 1000 Genomes Projects. Data for the canid genomes is available through NCBI.

## 6 Acknowledgments

We would like to thank Guhan Venkataraman for assistance in preparing the human dataset, Jan Sokol for sharing an initial implementation of the conditional random field used for comparison, and Trisha Singh for useful conversation on string kernel statistics. We also thank Eva Olafsdottir for her illustrations and invaluable contributions to visual design. This work was supported by the Chan Zuckerberg Biohub (CDB), National Library of Medicine (NLM) T15LM007033 (AGI), and the NIH Genome Sequencing Program 7U01HG009080.

## A Dataset compositions

We use 4 different simulated datasets: Latin American (AFR, EUR, NAT), 5-Ancestry (AFR, EUR, NAT, SAS, EAS), 7-ancestry (AFR, EUR, NAT, SAS, EAS, OCE, WAS), and Dev (AFR, EUR, NAT, SAS, EAS, OCE, WAS). Latin American, 5-Ancestry, and 7-Ancestry are used as benchmark datasets, and the Dev dataset is used for method development and hyper-parameter searches. Note that Dev dataset uses sequences from a different chromosome than the benchmarking datasets. Details of the numbers of single-ancestry individuals used for the simulation can be found in Table 1.

## B More on smoother data

The training set is split into two parts, one for training the base model and one for the smoother, to avoid distributional shift. Figures A1 and A2 show how the estimated probabilities have different distributions for the training data on the one hand and the independent validation dataset on the other. The dataset used is the dev dataset and the base module is XGBoost. It is clear that the estimated probabilities on the training data are sharper. That is, these probabilities tend to have much higher confidence (probability of prediction, maximum probability).

Furthermore, we argue that the smoothing module is not affected by the reduction in data, as it only needs a small fraction of the training data to perform properly as can be seen in Figure A3. The dataset is the same as above with the XGBoost smoothing module.

**Figure A1.**
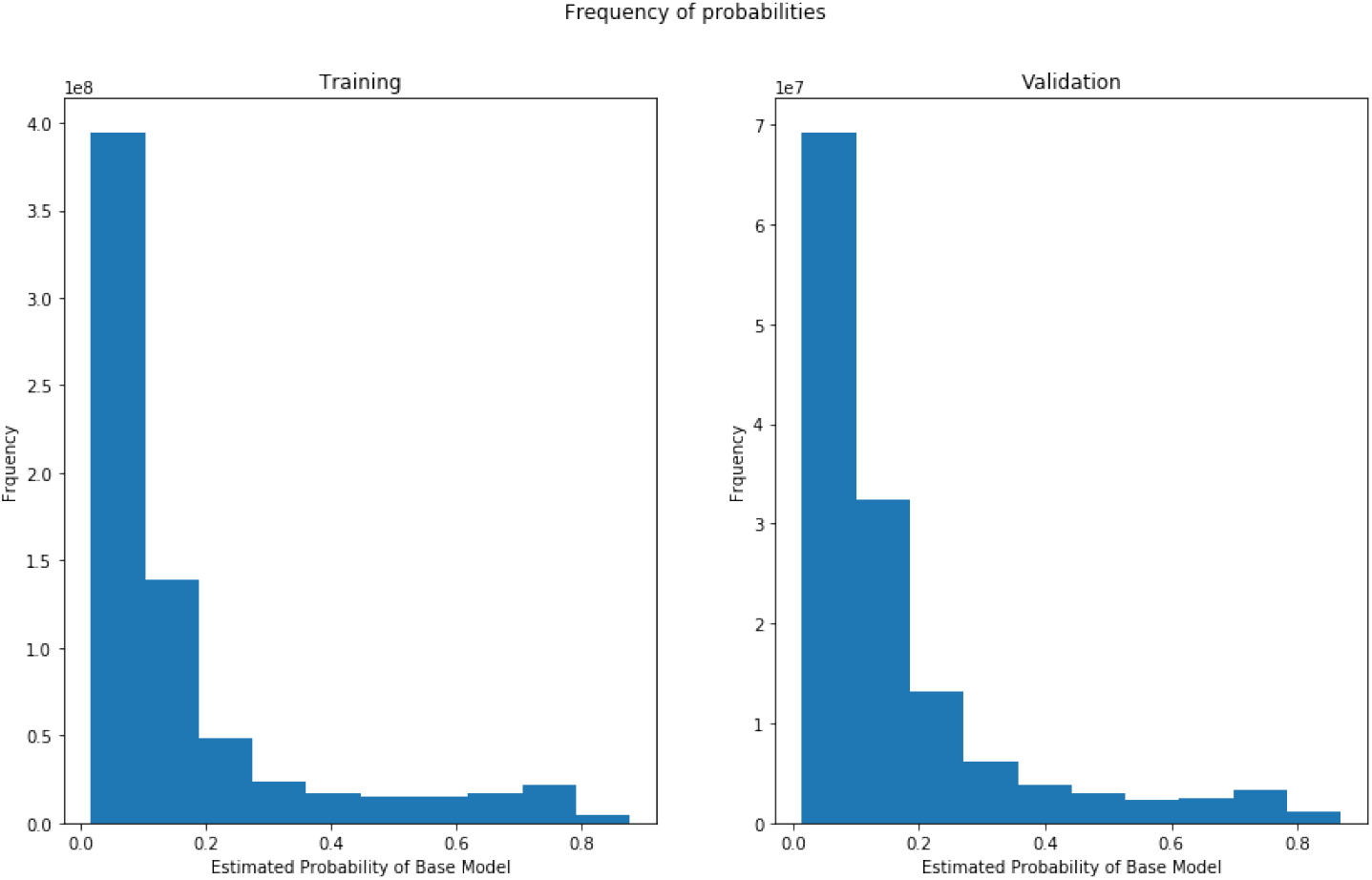
Different distributions of probability outputs for training individuals (left) and validation individuals (right).

**Figure A2.**
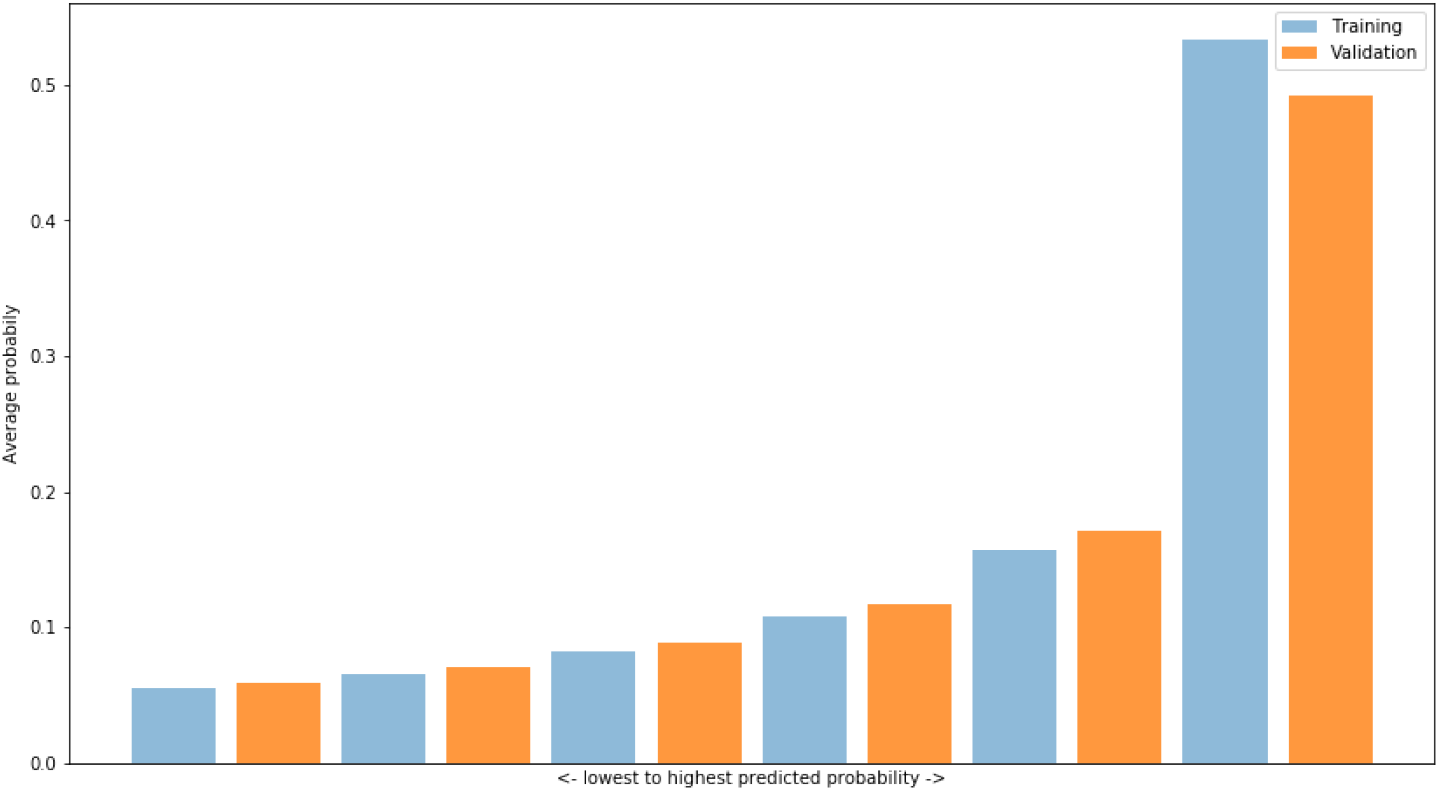
Average probabilities of ranked probability output.

**Figure A3.**
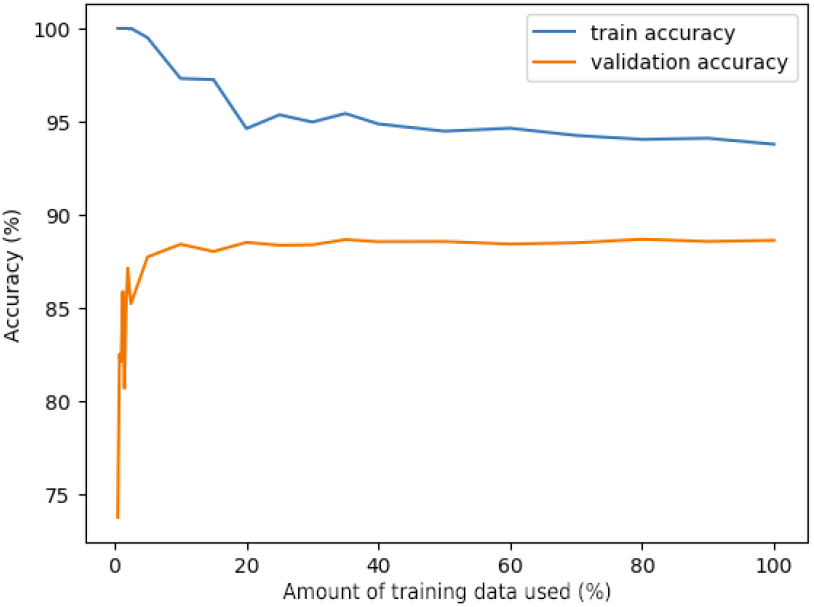
Smoother training saturation, eventually the smoother stops learning from additional data.

## C Empirical studies of calibration

For quantitative evaluation, we compare log-loss and calibration error^43^, and we find that both improve as a result of calibration. A comparison of the probabilities before and after calibration can be seen in the reliability plots in Figure 5. The figure shows the estimated probabilities and their respective true probabilities for two ancestries (West Asian and indigenous American). The mismatch between estimated and true probabilities is larger before calibration (dashed lines) than after (solid lines). For this dataset, the calibration error was 0.0257 before calibration and 0.0145 after. The log loss decreased from 0.3349 to 0.3249.

**Figure A4.**
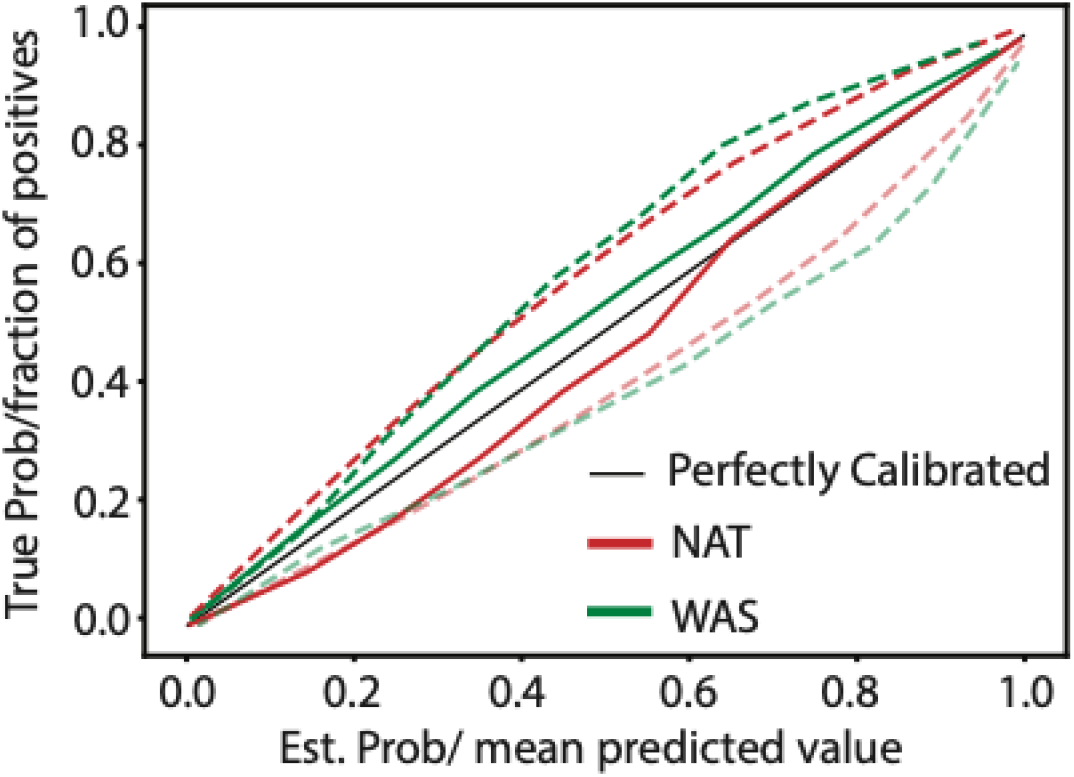
Reliability plot for before (dashed) and after (solid) calibration. Closer to the diagonal (perfectly calibrated, black) is better. The uncalibrated curves are mirrored across the diagonal to visually illustrate the centrality of the calibrated ones.

## D Benchmarking tables

We include a detailed benchmark of our different system’s configurations and competing methods on four different human datasets: Latin American, 5-Ancestry, and 7-Ancestry with whole-genome and genotyping array (SNPs used in UK Biobank Axiom array). Tables 2, 3, 4, 5, 6, 7 present the classification accuracy, log loss, base module classification accuracy, training time, inference time, and model size of each of the methods. Note that ELAI and Loter are only benchmarked with the 7-Ancestry dataset due to their extremely high computational requirements.

**Table 2.**
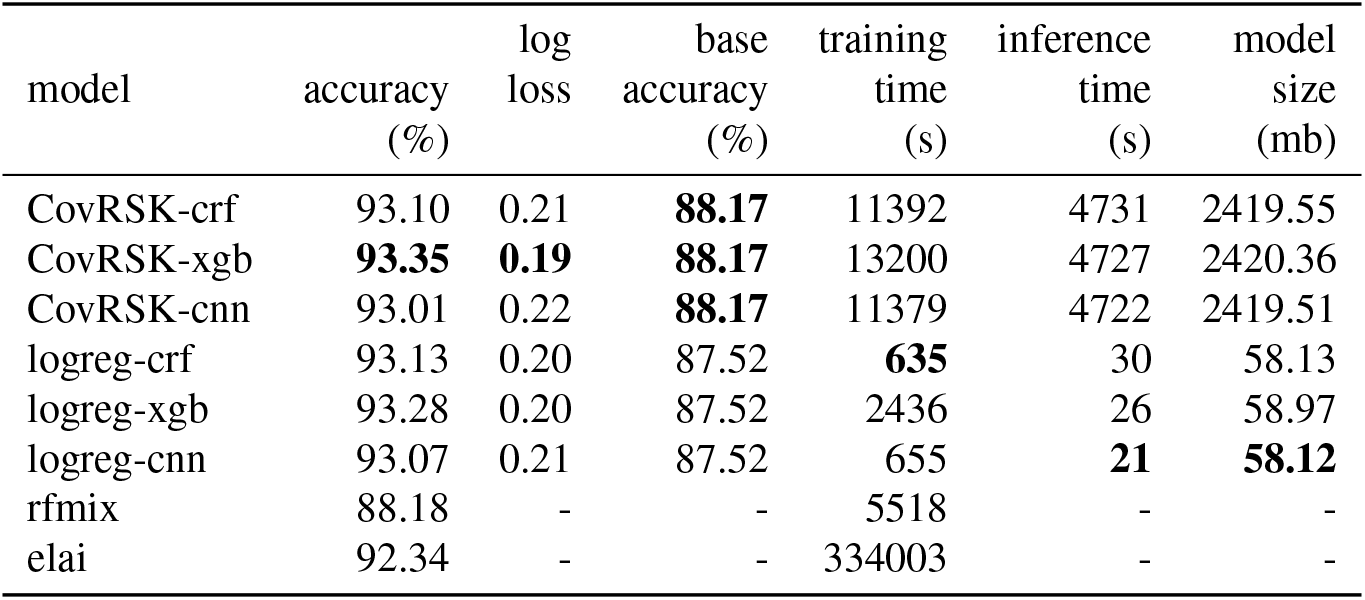
7-ancestry (whole genome)

| model | accuracy<br>(%) | log<br>loss | base<br>accuracy<br>(%) | training<br>time<br>(s) | inference<br>time<br>(s) | model<br>size<br>(mb) |
| --- | --- | --- | --- | --- | --- | --- |
| CovRSK-crf | 93.10 | 0.21 | **88.17** | 11392 | 4731 | 2419.55 |
| CovRSK-xgb | **93.35** | **0.19** | **88.17** | 13200 | 4727 | 2420.36 |
| CovRSK-cnn | 93.01 | 0.22 | **88.17** | 11379 | 4722 | 2419.51 |
| logreg-crf | 93.13 | 0.20 | 87.52 | **635** | 30 | 58.13 |
| logreg-xgb | 93.28 | 0.20 | 87.52 | 2436 | 26 | 58.97 |
| logreg-cnn | 93.07 | 0.21 | 87.52 | 655 | **21** | **58.12** |
| rfmix | 88.18 | - | - | 5518 | - | - |
| elai | 92.34 | - | - | 334003 | - | - |

**Table 3.** 5-ancestry (whole genome).

| model | accuracy<br>(%) | log<br>loss | base<br>accuracy<br>(%) | training<br>time<br>(s) | inference<br>time<br>(s) | model<br>size<br>(mb) |
| --- | --- | --- | --- | --- | --- | --- |
| CovRSK-crf | 95.16 | 0.14 | **91.50** | 11519 | 5004 | 2257.62 |
| CovRSK-xgb | **95.40** | 0.13 | **91.50** | 12608 | 5001 | 2258.22 |
| CovRSK-cnn | 95.07 | 0.14 | **91.50** | 11549 | 4997 | 2257.61 |
| logreg-crf | 95.19 | 0.14 | 91.40 | **433** | 23 | **41.57** |
| logreg-xgb | **95.40** | **0.12** | 91.40 | 1344 | 20 | 42.18 |
| logreg-cnn | 95.09 | 0.14 | 91.40 | 465 | **17** | **41.57** |
| rfmix | 90.76 | - | - | 4261 | - | - |

**Table 4.**
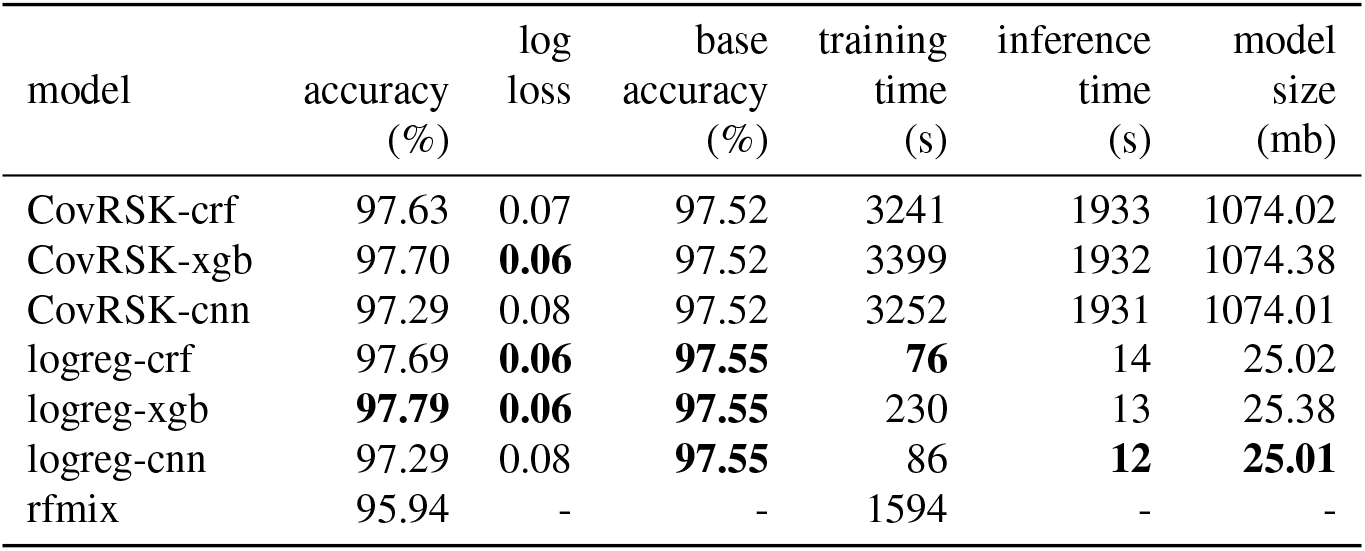
Latin American (whole genome).

| model | accuracy<br>(%) | log<br>loss | base<br>accuracy<br>(%) | training<br>time<br>(s) | inference<br>time<br>(s) | model<br>size<br>(mb) |
| --- | --- | --- | --- | --- | --- | --- |
| CovRSK-crf | 97.63 | 0.07 | 97.52 | 3241 | 1933 | 1074.02 |
| CovRSK-xgb | 97.70 | **0.06** | 97.52 | 3399 | 1932 | 1074.38 |
| CovRSK-cnn | 97.29 | 0.08 | 97.52 | 3252 | 1931 | 1074.01 |
| logreg-crf | 97.69 | **0.06** | **97.55** | **76** | 14 | 25.02 |
| logreg-xgb | **97.79** | **0.06** | **97.55** | 230 | 13 | 25.38 |
| logreg-cnn | 97.29 | 0.08 | **97.55** | 86 | **12** | **25.01** |
| rfmix | 95.94 | - | - | 1594 | - | - |

**Table 5.** 7-ancestry (array genotype).

| model | accuracy<br>(%) | log<br>loss | base<br>accuracy<br>(%) | training<br>time<br>(s) | inference<br>time<br>(s) | model<br>size<br>(mb) |
| --- | --- | --- | --- | --- | --- | --- |
| CovRSK-crf | 89.20 | **0.33** | **77.97** | 324 | 97 | 108.53 |
| CovRSK-xgb | **89.35** | **0.33** | **77.97** | 2470 | 93 | 109.35 |
| CovRSK-cnn | 89.07 | 0.35 | **77.97** | 324 | 87 | 108.50 |
| logreg-crf | 86.57 | 0.40 | 67.90 | 1402 | 11 | **2.06** |
| logreg-xgb | 86.01 | 0.42 | 67.90 | 3792 | 7 | 2.91 |
| logreg-cnn | 86.31 | 0.41 | 67.90 | 1555 | **1** | **2.06** |
| rfmix | 84.81 | - | - | **241** | - | - |
| elai | 88.32 | - | - | 37219 | - | - |
| loter | 83.86 | - | - | 8840 | - | - |

**Table 6.** 5-ancestry (array genotype).

| model | accuracy<br>(%) | log<br>loss | base<br>accuracy<br>(%) | training<br>time<br>(s) | inference<br>time<br>(s) | model<br>size<br>(mb) |
| --- | --- | --- | --- | --- | --- | --- |
| CovRSK-crf | 91.85 | 0.24 | **81.61** | 241 | 86 | 89.55 |
| CovRSK-xgb | **92.06** | **0.23** | **81.61** | 1307 | 83 | 90.15 |
| CovRSK-cnn | 91.72 | 0.24 | **81.61** | 272 | 79 | 89.54 |
| logreg-crf | 90.08 | 0.29 | 74.46 | **37** | 15 | **1.54** |
| logreg-xgb | 89.99 | 0.29 | 74.46 | 887 | 13 | 2.15 |
| logreg-cnn | 90.00 | 0.29 | 74.46 | 70 | **9** | **1.54** |
| rfmix | 90.28 | - | - | 196 | - | - |

**Table 7.** Latin American (array genotype).

| model | accuracy<br>(%) | log<br>loss | base<br>accuracy<br>(%) | training<br>time<br>(s) | inference<br>time<br>(s) | model<br>size<br>(mb) |
| --- | --- | --- | --- | --- | --- | --- |
| CovRSK-crf | 96.55 | 0.10 | **93.86** | 67 | 32 | 36.78 |
| CovRSK-xgb | **96.67** | **0.09** | **93.86** | 233 | 31 | 37.14 |
| CovRSK-cnn | 96.36 | 0.11 | **93.86** | 79 | 29 | 36.77 |
| logreg-crf | 96.23 | 0.11 | 91.68 | **12** | 8 | 1.02 |
| logreg-xgb | 96.30 | 0.10 | 91.68 | 168 | 7 | 1.37 |
| logreg-cnn | 95.90 | 0.12 | 91.68 | 23 | **6** | **1.00** |
| rfmix | 95.58 | - | - | 82 | - | - |

## E CovRSK theory and extended experiments

### E.1 String kernel features as Binomial random variables

Assuming that the input features (SNPs in this case) are binary, as well as independent and identically distributed,

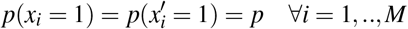

Then the probability that a given feature is the same for two individuals is,

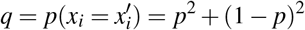

The event that a given subsequence of length 1 is equal for two individuals is clearly Bernoulli distributed and thus the number of subsequences of length 1 is binomial, *S*_1_ ~ Binomial(M,q) where M is sequence length. The expected value and variance is then given by,

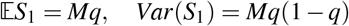

Now, given our assumptions, two sequences of length *k* will be equal with probability *q* given that the first *k* – 1 elements are equal and thus,

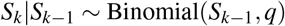

with expected value and variance,

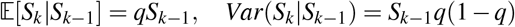

and therefore,

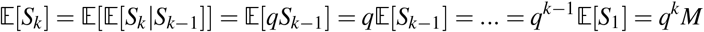

and

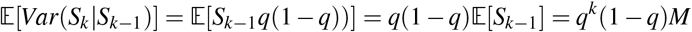

We visualize the last expression (normalized by the sequence length) in Figure A5.

### E.2 String kernel feature covariance

By modelling the string kernel features as binomials, the expected value is given by,

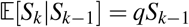

and thus the covariance can be computed using the law of total expectation,

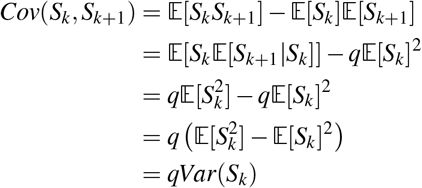

**Figure A5.**
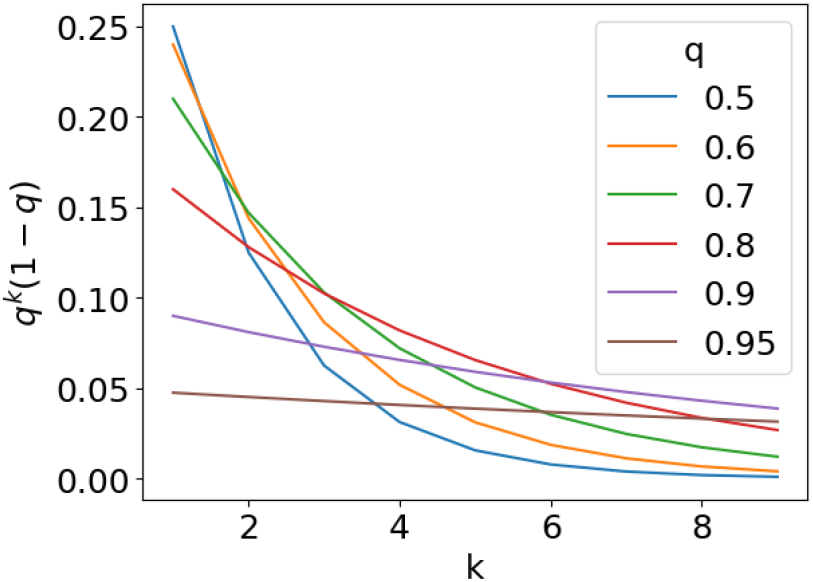
Expectation of the variance of string kernel features, given the previous feature, decreases exponentially in k.

Using the recurrence in E.1, this is easily extended to obtain the covariance between two arbitrary string kernel features,

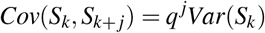

### E.3 String kernel feature correlation

By modelling the string kernel features as binomials, the expected value is given by,

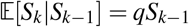

and as showed in E.2, the covariance is given by,

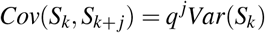

A lower bound on the correlation can then be computed using the law of total variance,

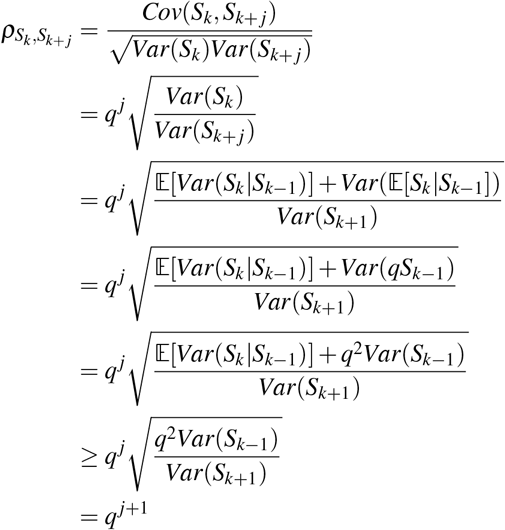

**Figure A6.**
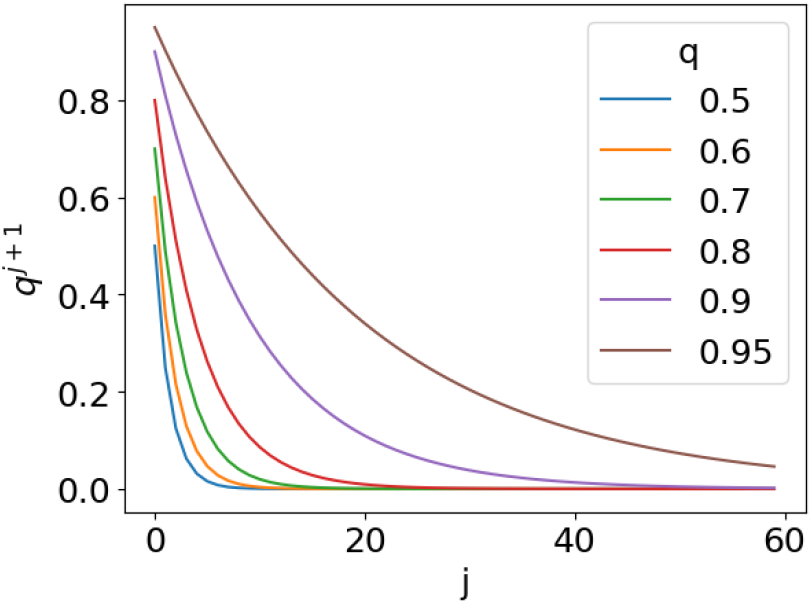
Lower bound on string kernel feature correlation decays exponentially with their distance.

which is visualized in Figure A6.

### E.4 String kernel thought experiment

**Figure A7.**
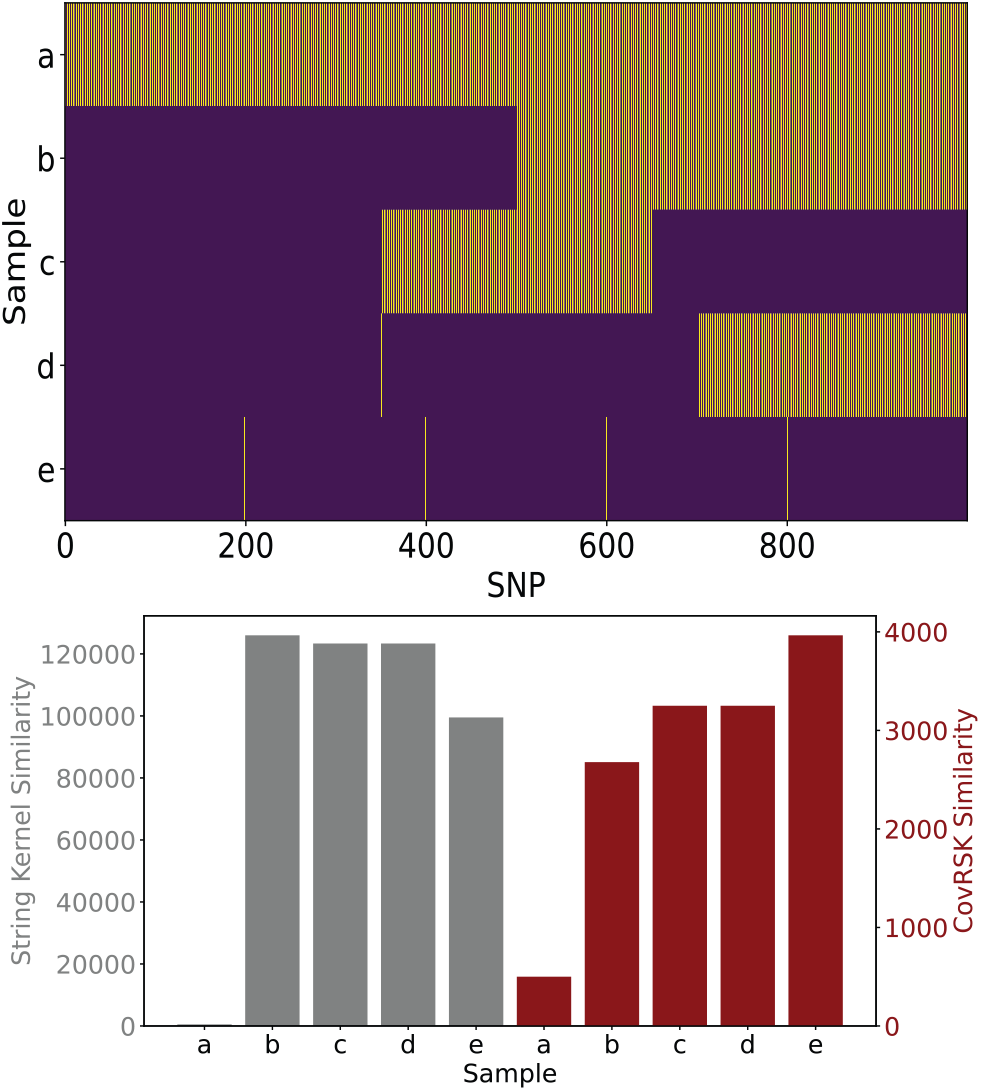
CovRSK thought experiment. Above, we have created samples where purple represents matching SNPs and yellow mismatches. Below, we show the kernel similarity for both the string kernel (gray) and CovRSK with *α* = 0.9 and *β* = 0.5 (red).

To provide more insight, we demonstrate a key difference between CovRSK and the standard string kernel using an example, which is visualized in Figure A7. Sample *b* shares only 50% of its SNPs with the example reference sample (the first half matches the reference exactly), while sample *e* shares 99.5% of its SNPs with the reference sample (but has shorter contiguous subsequences of exact match than the first half of sample *b*). Sample *e* has the highest CovRSK similarity to the reference sample, but the string kernel ranks sample *b* to be the most similar. This is due to the accumulation of the correlated string kernel features. As the contiguous sequence match gets longer, the number of total k-mers matching grows by the length of the contiguous match (one for each length, from one to the full contiguous match length) meaning that the match’s overall contribution to the similarity score grows as O(*n*^2^) with its length. As a result of this nonlinearity, the long contiguous sequence match of sample *b* contributes more than the sum of the many - but shorter - sequence matches of sample *e*. This characteristic of the string kernel need not be bad and might in some cases even be desired. However, given the stochastic nature of genetic inheritance and potential for noisy sequencing or genotyping, one cannot expect sequences of related individuals in a population to share the exact same contiguous sequence, even though the two individuals may have inherited the contiguous sequence identically-by-descent from a common ancestor. It is more likely that the sequences of the two individuals’ genomes stemming from the same population will have very low variance in general. Therefore, the stochasticity of the contiguous length of the shared sequence along with the quadratically increasing influence on the string kernel of segment length could potentially lead to overfitting to the training samples. In general, the CovRSK vies with the standard string kernel for best accuracy, beating all other methods with the top spot trading (narrowly) between them depending on the smoother used; however, in all cases the CovRSK is far faster than the regular string kernel.

### E.5 Empirical results

In Figure A8 we see a comparison between CovRSK and the string kernel similar to the example above, but this time with real samples. Again the samples are colored such that purple represents SNPs matching the reference sample and yellow denotes mismatches. The only sample actually sharing ancestry with the reference sample is sample four. We notice how CovRSK correctly returns that sample as the most similar, while the string kernel incorrectly scores sample seven higher.

**Figure A8.**
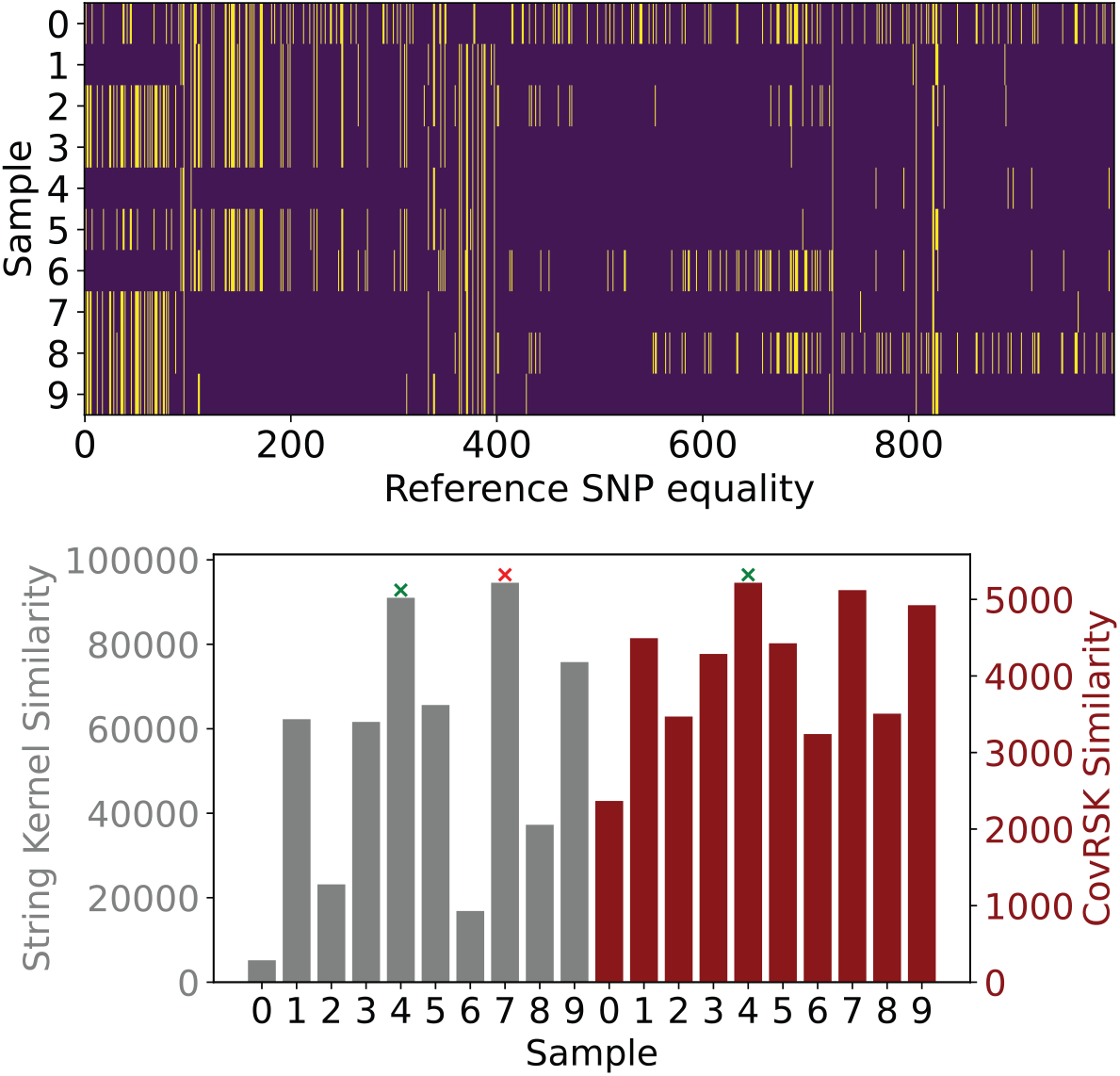
Empirical example. Above, we have random samples where purple represents shared SNP sequences with a random reference sample, and yellow denotes differences. The populations are the same as in the seven ancestry dataset and only sample four has the same ancestry as the reference. Below we show kernel similarity for both the string kernel (gray) and CovRSK with *α* = 0.9 and *β* = 0.5 (red).

To provide insight into the CovRSK sampling algorithm, an instance on a sequence of length 300 is visualized in Figure A9. The blue solid line represents the probability of sampling a *k* SNP sequence given the sampling history. The dashed line represents the decaying probability without covariance reduction. As the subsequences get larger, the sampling probability decreases in general. When a *k* length sequence is sampled, the probability of sampling the next higher *k* drops and slowly rises as the distance increases.

**Figure A9.**
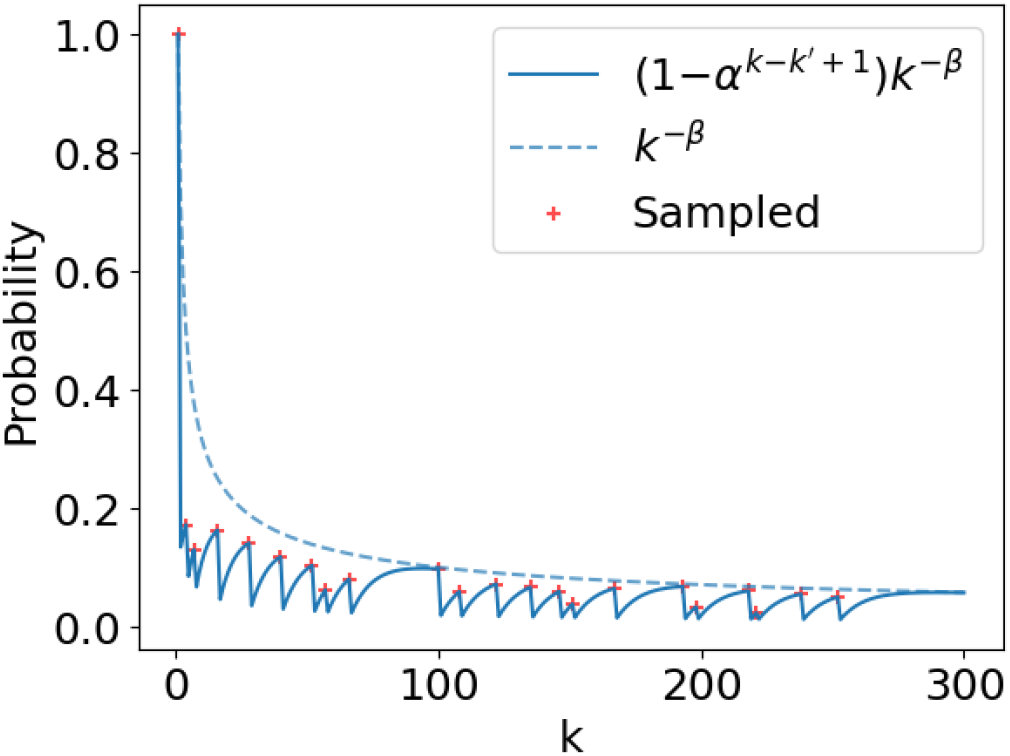
An instance of CovRSK feature samples and their sample probability with *α* = 0.9 and *β* = 0.5.

## F Algorithms for string kernel computation

In this section, we list various string kernel algorithms mentioned in section 2.2. In Algorithm 5 we have a detailed, lower level description of the string kernel computation with triangular numbers. In Algorithm 6 we then have the dynamic programming (DP) version and finally, in Algorithm 7, we list the DP version that is customized for the CovRSK.

### Algorithm 5: Detailed algorithm for string kernel with triangular numbers

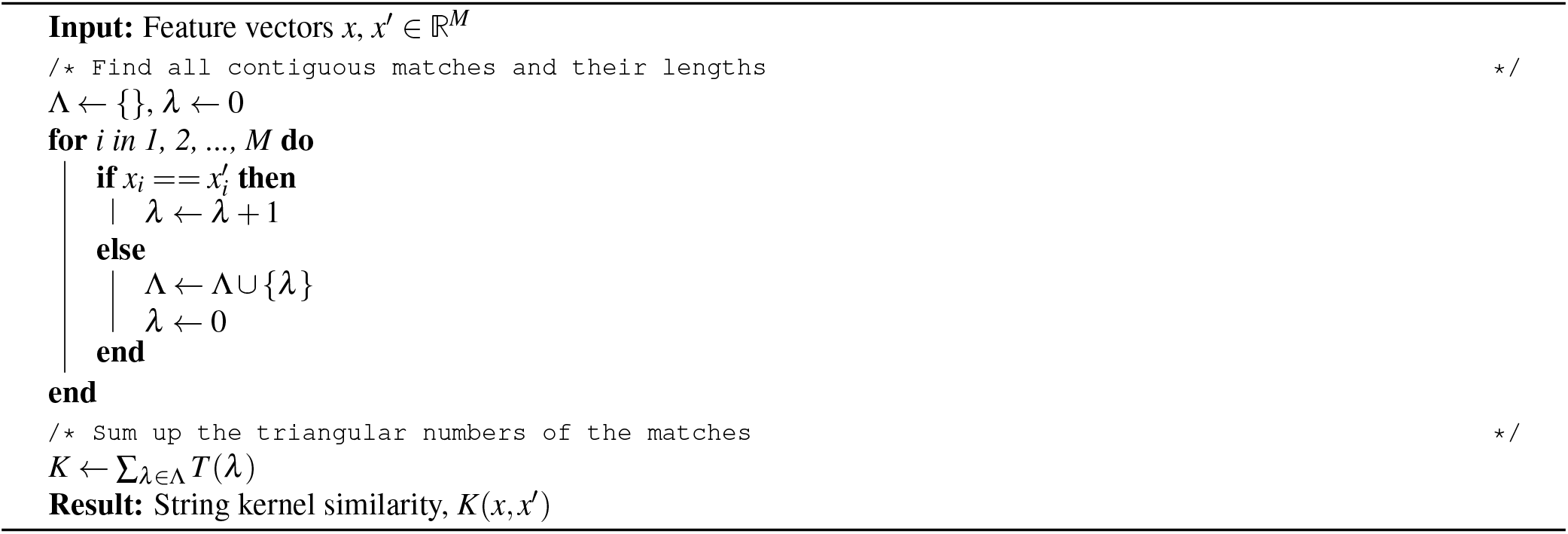

### Algorithm 6: String kernel with triangular numbers and DP

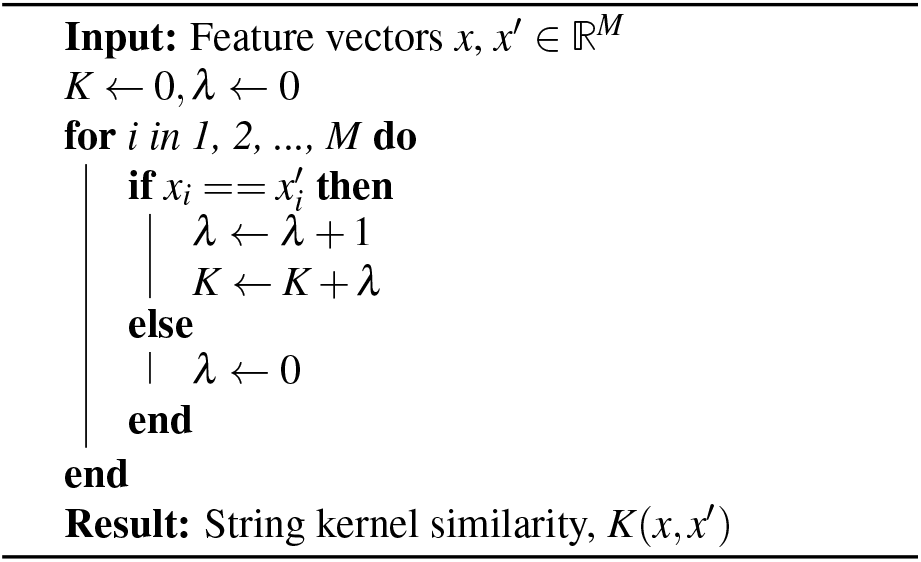

### Algorithm 7: DP version of CovRSK

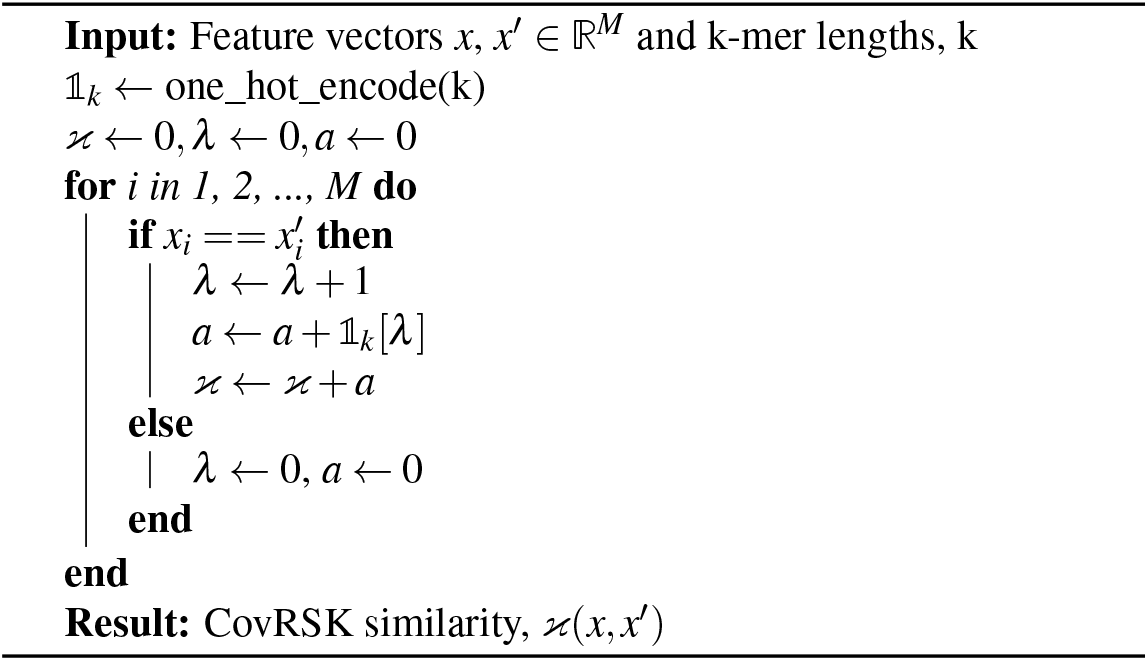

## G Error analysis

While model predictions are important, it is also necessary to understand signs of uncertainty in the predicted estimates. In figure A10, we show how the model is more likely to make mistakes near ancestry switch points (break points). This explains in part the decay of accuracy as generation time since admixture increases. With higher generation time, we have more frequent breakpoints leading to a more difficult problem.

**Figure A10.**
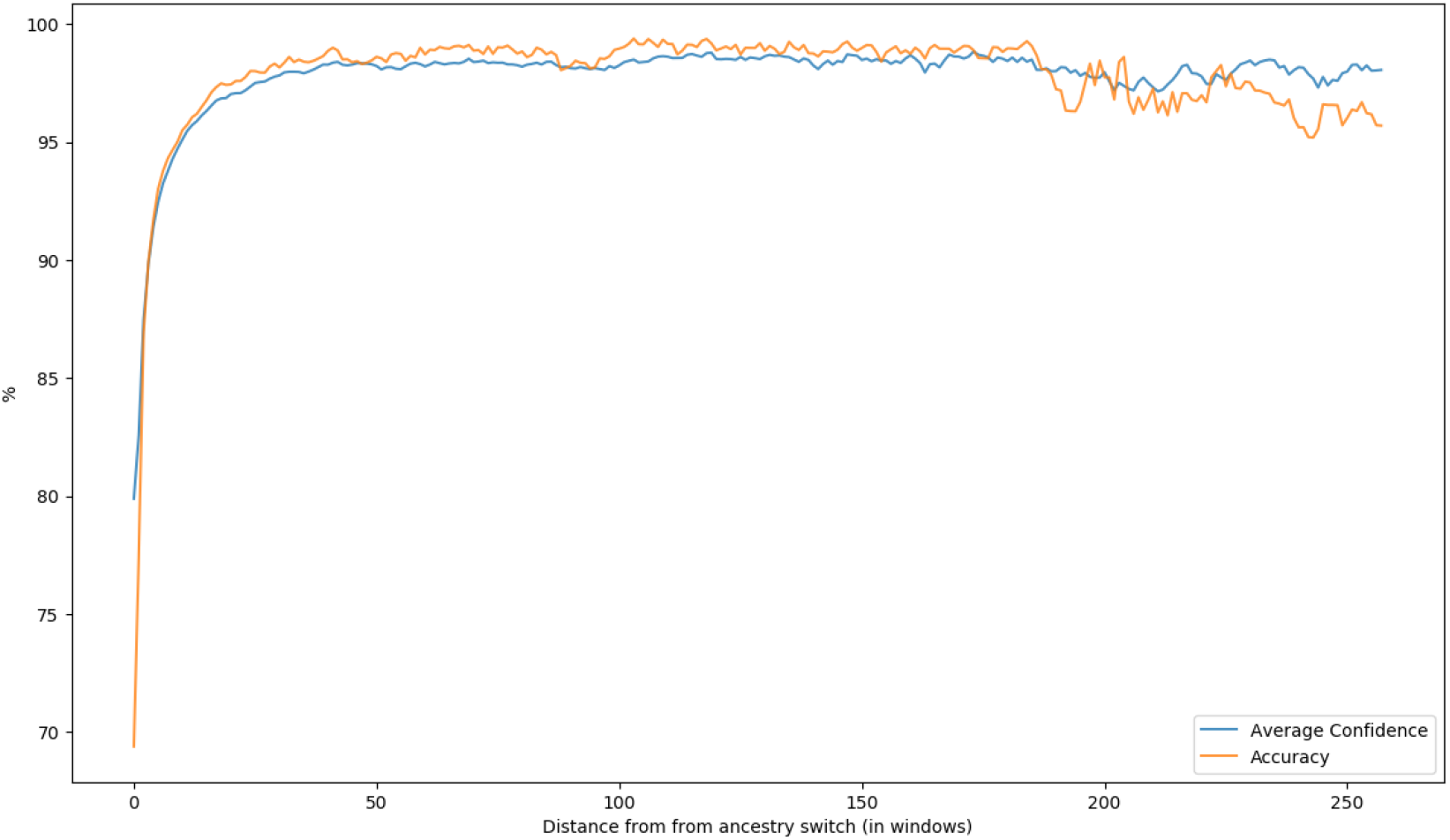
Validation accuracy as a function of distance from ancestry switch (in windows).

In Figures A11 and A12 we plot confusion matrices for Gnomix and RFMix respectively on generation 64 validation data in the dev dataset. It’s clear that some population pairs present a harder classification task than others, for instance the neighboring European (EUR) and West Asian (WAS) ancestries. The two methods seem to make similar mistakes, but Gnomix has fewer of them.

**Figure A11.**
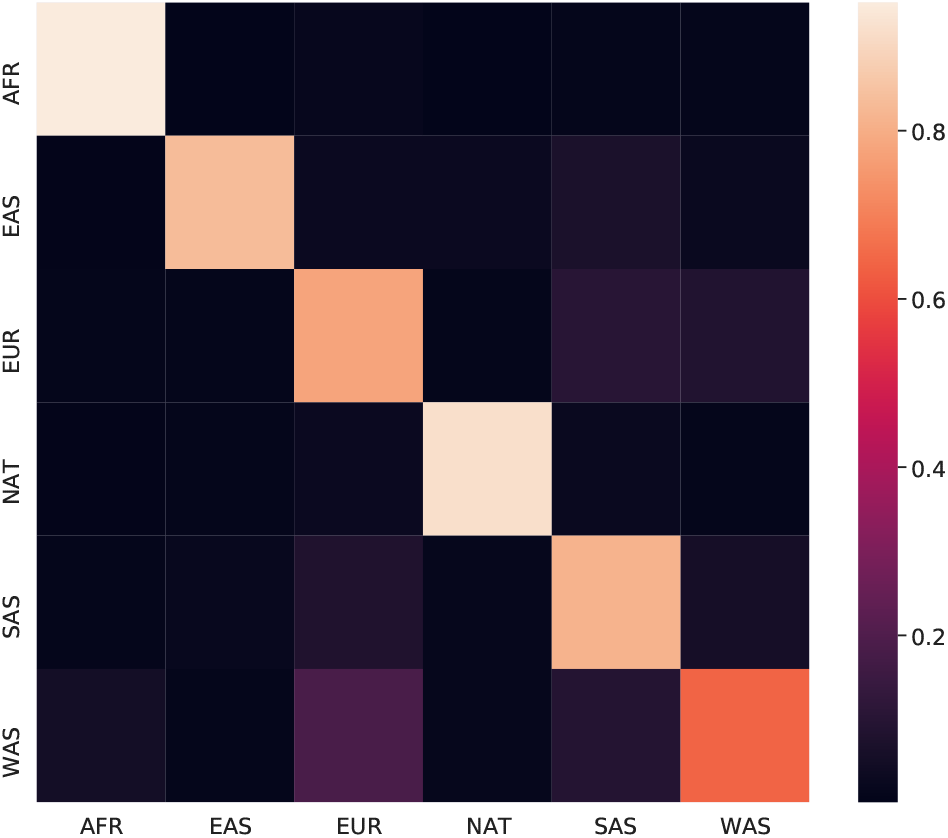
Gnomix normalized confusion matrix on the dev dataset.

**Figure A12.**
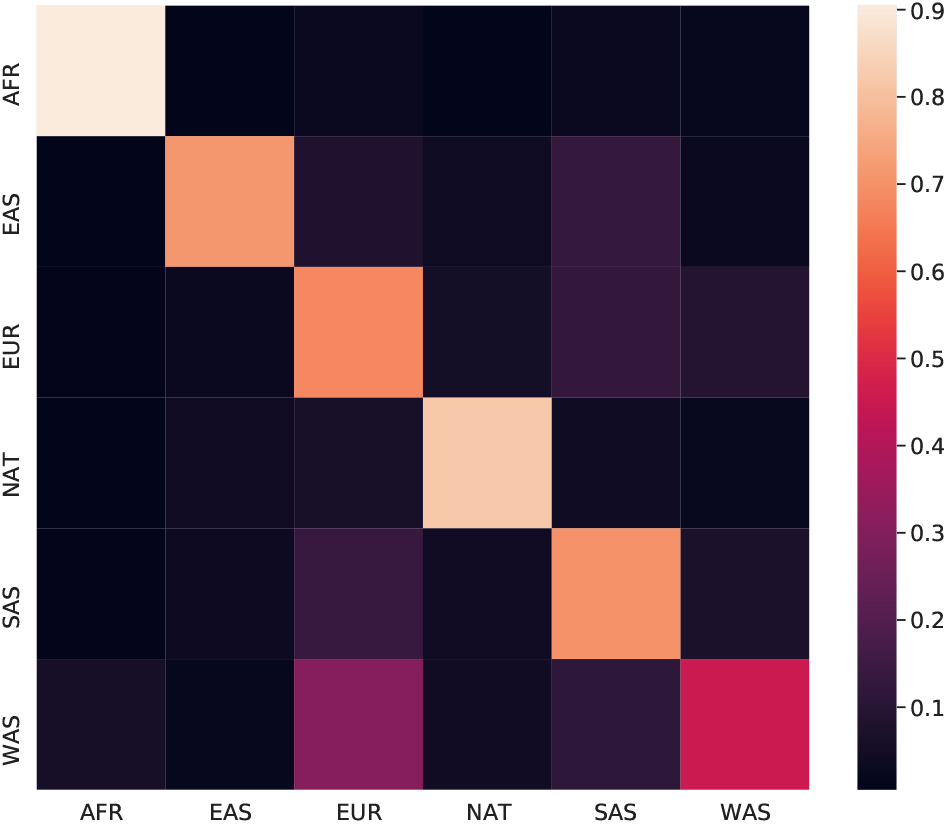
RFMix normalized confusion matrix on the dev dataset.

## H More on phasing error correction

We use a dataset of simulated admixed individuals with known local ancestry labels to create a dataset containing phasing errors. Since each segment from the single ancestry dataset is correctly phased up to its ancestry, we can create an instance of correctly ancestry phased individual by collapsing the simulated admixed indivduals created from them to unphased genotypes and then applying standard phasing software (here Beagle 5.1). This procedure is described in more detail in Algorithm 8 and produces the dataset for evaluation that we refer to as the Latin American phasing dataset. The simulated individuals model modern Latin American individuals with eight generations since the onset of admixture between indigenous American, African and European ancestries.

### Algorithm 8: Simulating data containing phasing errors

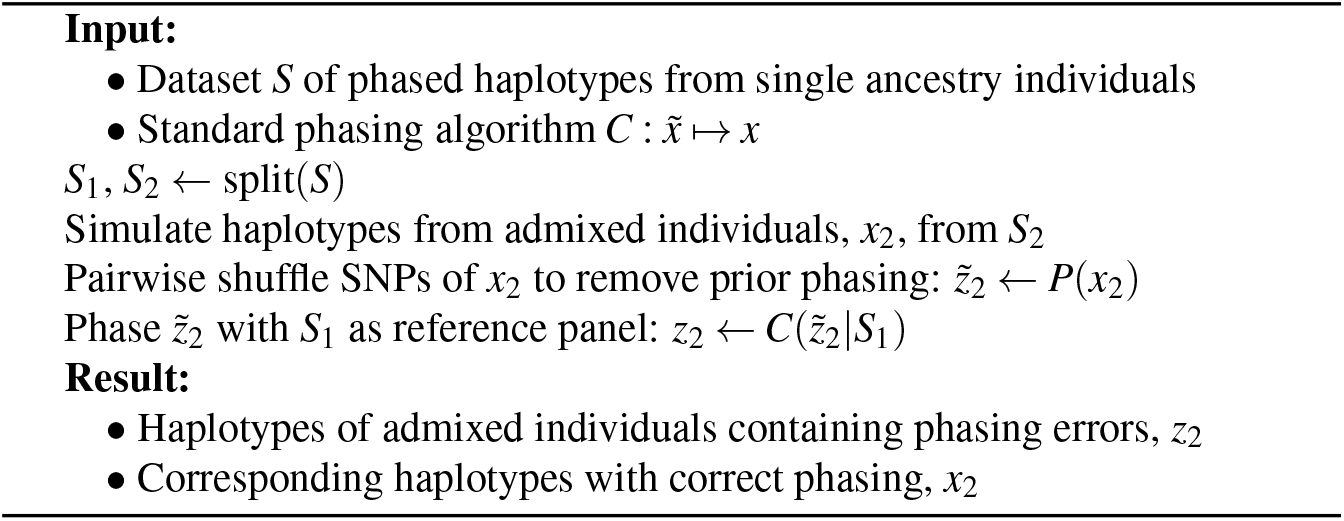

**Table 8.**
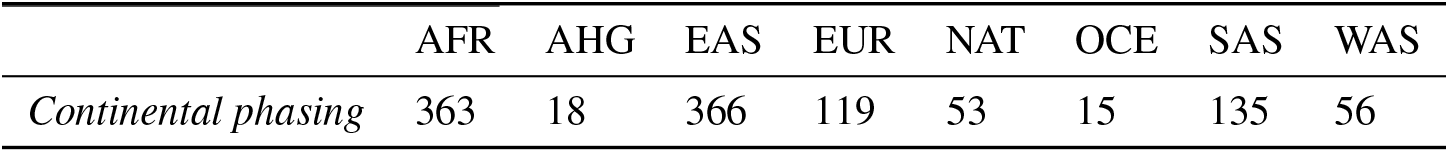
Single ancestry individual composition in the simulated datasets.

|  | AFR | AHG | EAS | EUR | NAT | OCE | SAS | WAS |
| --- | --- | --- | --- | --- | --- | --- | --- | --- |
| *Continental phasing* | 363 | 18 | 366 | 119 | 53 | 15 | 135 | 56 |

**Table 9.** Phasing and phasing error correction run-times in seconds with relative speed-up compared with RFMix.

|  | Latin American | continental |
| --- | --- | --- |
| Beagle 5.1 | 316 | 800 |
| RFMix | 2,252 s (1×) | 172,800 s (1×) |
| Gnofix | **17 s** (132.5×) | **98 s** (1,763.3×) |

RFMix’s phasing error correction algorithm is hard to separately time, since its phase correction and ancestry estimation are intertwined. A lower bound on the correction time can be estimated by measuring RFMix’s run-time for performing LAI without phase correction and then subtracting this from its time run-time for performing combined LAI and phase correction. When the number of ancestries becomes sizeable (e.g. more than five) the run-time is hard to exhaustively measure for many individuals. As we can see the lower bound on the phase correction run-time for RFMix increases superlinearly with the number of ancestries. Indeed, at just over 10 ancestries the phase correction in RFMix becomes infeasible for any sizeable number of query individuals.

**Figure A13.**
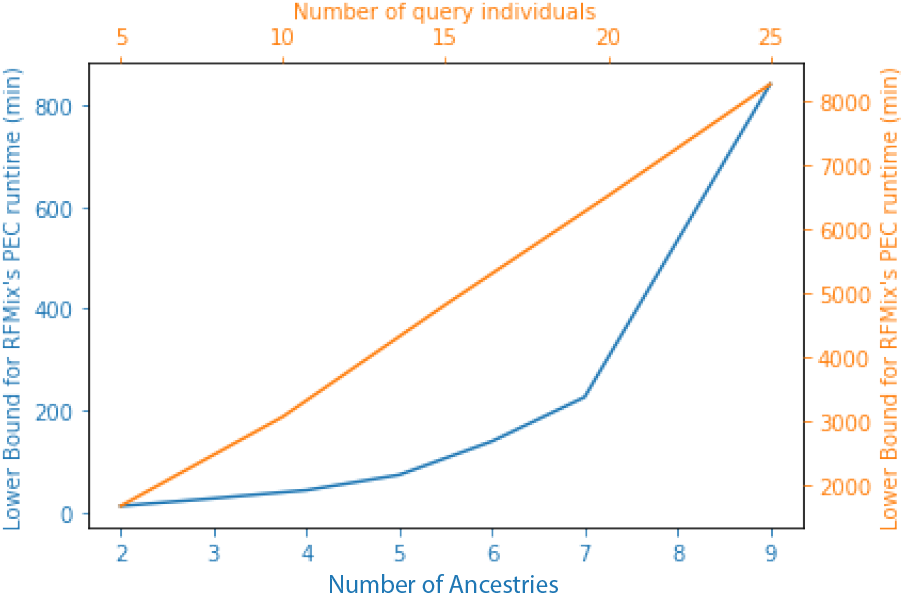
RFMix’s run-time for a given number of query individuals (orange) and different ancestries (blue). In the first case, the reference panel size and the number of ancestries was kept constant at 350 and three respectively. In the latter, the reference panel size and number of query individuals was kept constant at 240 and one respectively.

